# Probing intracellular physical environments by rotational and translational single-particle tracking

**DOI:** 10.64898/2026.08.14.744592

**Authors:** Dimitri Schumacher, Martin D. Baaske, Wenxin Zhang, Biswajit Pradhan, Delong Li, Thorsten Feichtner, Florian Wilfling, Eugene Kim

## Abstract

Single-particle tracking is widely used to probe nanoscale dynamics in biological systems, yet most approaches rely exclusively on translational motion, overlooking rotational dynamics that offer complementary information about the local physical environment. Here, we present a simultaneous rotational and translational single-particle tracking approach using a vortex-engineered point spread function in a single detection channel. We validate this approach with static and freely diffusing nanorods, demonstrating accurate orientation recovery and quantitative agreement with theoretical predictions of rotational diffusion. Using a biomimetic lipid bilayer system, we show that translational and rotational diffusion exhibit distinct sensitivities to environmental perturbations, confirming that these two modalities capture complementary local environment information. Applying this framework to living HeLa cells, we show that combined translational and rotational diffusion signatures define distinct biophysical fingerprints of cytoplasmic and endocytic compartments and reveal compartment-specific responses to metabolic perturbation. Finally, time-resolved analysis of individual endocytic compartments uncovers dynamic changes in the local physical environment that are inaccessible to conventional translational tracking. By coupling translational and rotational readouts, this framework opens a new dimension for probing the physical organization and dynamics of living systems at the nanoscale.

## Introduction

Single-particle tracking has become an indispensable approach for probing the dynamic behavior of biological systems [^1,2^]. By following the trajectories of individual molecules, nanoparticles, and intracellular structures, measurements of translational motion have provided fundamental insights into intracellular transport, molecular interactions, and the physical properties of cellular environments. However, translational motion represents only one aspect of particle dynamics. Anisotropic particles also undergo rotational motion, which is governed by distinct physical principles and can provide complementary information about the local physical environment that cannot be inferred from positional tracking alone [^3–6]^. Simultaneous measurements of translational and rotational dynamics therefore provide complementary biophysical readouts that enable a more comprehensive characterization of intracellular environments.

Several optical approaches have been developed to measure the orientation and rotational dynamics of individual fluorophores, primarily using polarized fluorescence or engineered point spread functions [^7–13]^. However, fluorescence-based approaches remain fundamentally limited by finite photon budgets and photobleaching, limiting observation times. Moreover, the rapid rotational diffusion of fluorophores often necessitates measurements of static orientations or the use of immobilized samples to avoid orientation averaging during image acquisition, limiting the applicability of these approaches for tracking rotational dynamics in living cells. Consequently, simultaneous long-term, high-speed measurements of translational and rotational dynamics in living cells remain challenging.

Anisotropic plasmonic nanoparticles, such as gold nanorods (AuNRs), provide an attractive alternative for orientation-resolved imaging [^5,14–16]^. Their localized surface plasmon resonance generates exceptionally strong and spectrally tunable scattering signals, while the elastic nature of optical scattering renders them immune to photobleaching, enabling prolonged observation [^17^]. In addition, their larger hydrodynamic dimensions slow rotational diffusion into timescales that can be readily resolved with high-speed cameras [^18^]. Together with their anisotropic geometry, which produces orientation-dependent scattering [^19^], these properties make gold nanorods an attractive platform for long-term simultaneous measurements of translational and rotational dynamics in living cells.

Here, we present a scattering-based microscopy approach for simultaneous translational and rotational tracking of individual gold nanorods using a vortex-engineered point spread function in a single detection channel. We first validate this approach in controlled model systems, establishing that translational and rotational diffusion report on distinct physical properties of the local environment. We then apply this framework to living cells to ask whether these complementary observables can resolve physical differences between intracellular compartments, capture their remodelling during physiological perturbation, and reveal dynamic changes that remain hidden to conventional positional tracking.

## Results

### Vortex-encoded orientation tracking of gold nanorods

To enable simultaneous measurements of nanoparticle position and orientation, we engineered the point spread function (PSF) using a vortex-phase plate (VPP, m=1) positioned in the infinity-conjugated plane of the microscope (Fig. 1A, Fig. S1). This configuration generates an orientation-encoding vortex-PSF (VPSF), thereby encoding both in-plane (ϕ) and out of plane (θ) angles of individual AuNRs within a single detection channel (Fig. 1B,C). A beam-stop placed in the focal plane of the reflected excitation beam creates a near-darkfield configuration that suppresses background and enhances the contrast of AuNR scattering, similar to strategies used in interferometric scattering microscopy [^20,21^].

**Figure 1.**
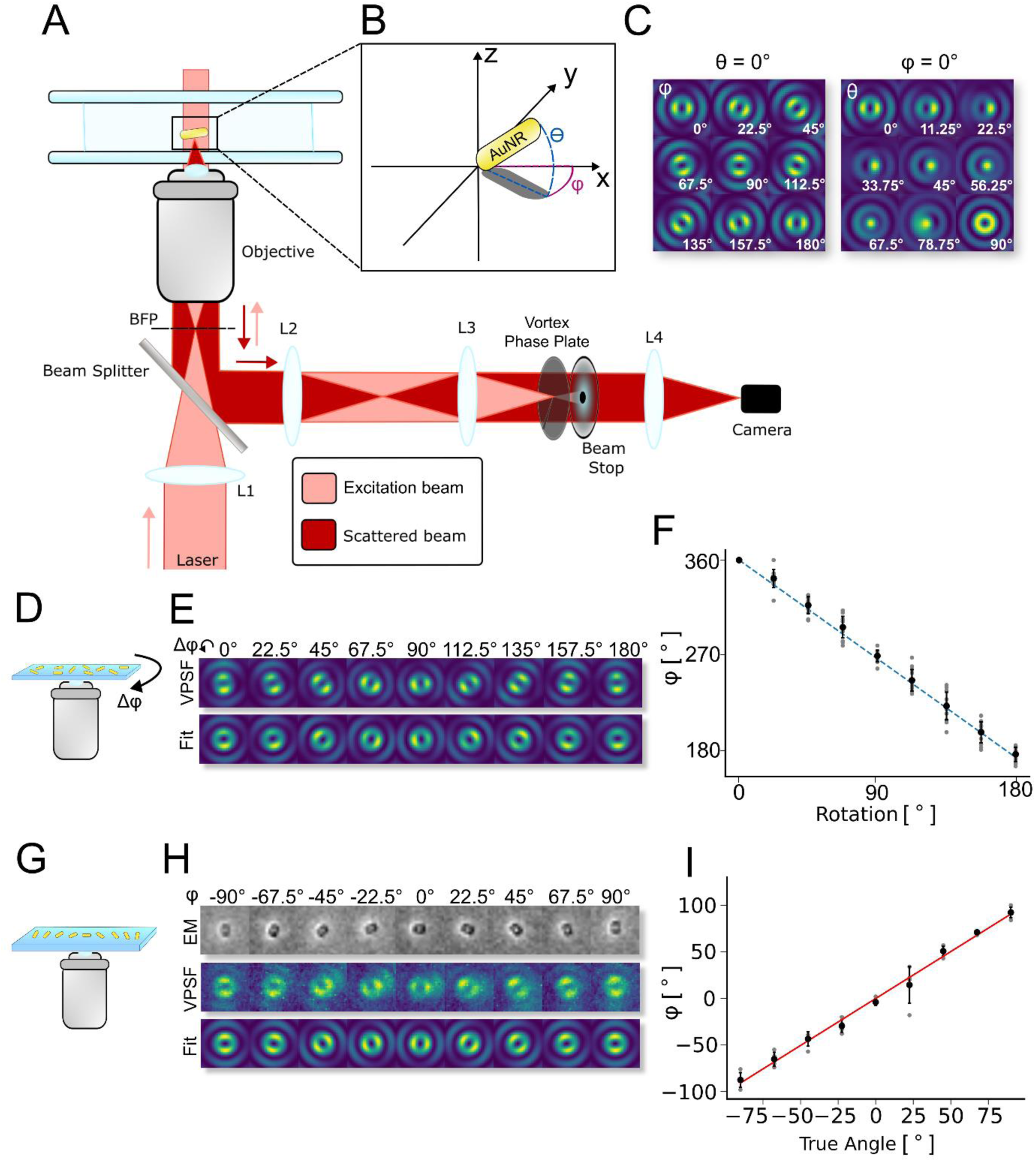
| Validation of vortex point spread function (VPSF)-based orientation tracking. **(A)** Schematic of the microscope used for simultaneous translational and rotational tracking of AuNRs. **(B)** Schematic showing the AuNR orientation angles, azimuthal angle (ϕ) and polar angle (θ). **(C)** Simulated VPSFs for representative rotations in ϕ (left) and θ (right). **(D)** Schematic of manually rotated AuNRs immobilized on a glass coverslip. **(E)** Representative VPSFs (top) and corresponding fits (bottom) during manual rotation of an AuNR through 180° in 22.5° increments in ϕ. **(F)** Measured relative ϕ rotation of manually rotated AuNRs on a glass coverslip. **(G)** Schematic of a nanofabricated AuNR grid containing AuNRs with predefined ϕ orientations spanning 180° in 22.5° increments. **(H)** Electron microscopy images of nanofabricated AuNRs (top) together with the corresponding VPSFs acquired in luminescence mode (middle) and the fitted orientations (bottom). **(I)** Measured versus nominal ϕ orientation of the nanofabricated AuNR grid. Linear regression with a slope of 1.01.

To determine particle orientations from raw image data, we implemented a brute-force-fitting algorithm that compares experimentally recorded VPSFs to a library of vectorially simulated VPSFs spanning all possible angular combinations with 1° resolution (see Supplementary Information). This approach enables frame-by-frame reconstruction of particle orientations and thus rotational trajectories.

We first benchmarked orientation recovery at the single-particle level using immobilized AuNRs mounted on a rotation stage. AuNRs were imaged in scattering mode while being rotated in defined 22.5° increments over 180° (Fig. 1D,E). The reconstructed in-plane (ϕ) angles closely followed the imposed rotations (Fig. 1F). Throughout the experiment, out-of-plane angles (θ) remained close to zero (θ_median_ = 5°; Fig. S2A), consistent with nanorods lying nearly flat on the substrate and confirming reliable three-dimensional orientation recovery.

To assess orientation precision and potential systematic bias, we analyzed randomly distributed immobilized AuNRs. The reconstructed in-plane angles exhibited an approximately uniform distribution, indicating the absence of angular bias (Fig. S2C-E). Frame-to-frame angular fluctuations yielded a mean standard deviation of approximately 5° for ϕ and 2.5° for θ (Fig. S2B). In addition, z-stack measurements over ±250 nm confirmed stable orientation recovery throughout the relevant focal volume, with maximal uncertainties of Δϕ ≈ ±10° and Δθ ≈ ±5° at the largest defocus distances examined (Fig. S2F-H).

Finally, we validated the absolute orientation accuracy using nanofabricated AuNR arrays with predefined orientations spanning 22.5° increments over 180° (Fig. 1G-I). The designed nanorod orientations were first verified by electron microscopy before optical measurements (Fig. S2I-J). VPSF imaging performed in photoluminescence mode (Fig. S2K) yielded reconstructed in-plane angles that closely matched the designed pattern, with a linear fit slope of 1.01 (Fig. 1I). As expected, reconstructed out-of-plane angles remained near zero (Fig. S2J).

Together, these experiments establish accurate, unbiased, and robust orientation recovery, providing the foundation for quantitative simultaneous measurements of translational and rotational dynamics.

### Quantitative measurements of rotational diffusion

Having established accurate orientation recovery, we next evaluated whether the platform could quantitatively resolve rotational dynamics under controlled conditions. Freely diffusing AuNRs were first examined in glycerol–water mixtures of varying viscosities. Representative snapshots of diffusing nanorods (Fig. 2A,B) and reconstructed trajectories (Fig. 2C) revealed continuous, stochastic rotational motion, with particles exploring a broad range of both in-plane (ϕ) and out-of-plane (θ) orientations (Fig. S3A-B). Rotational diffusion coefficients (D_r_) were extracted via mean square angular displacement (MSAD) analysis [^22,23^] (Fig. 2D):

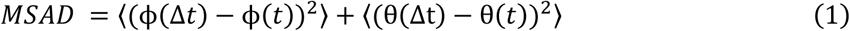

**Figure 2.**
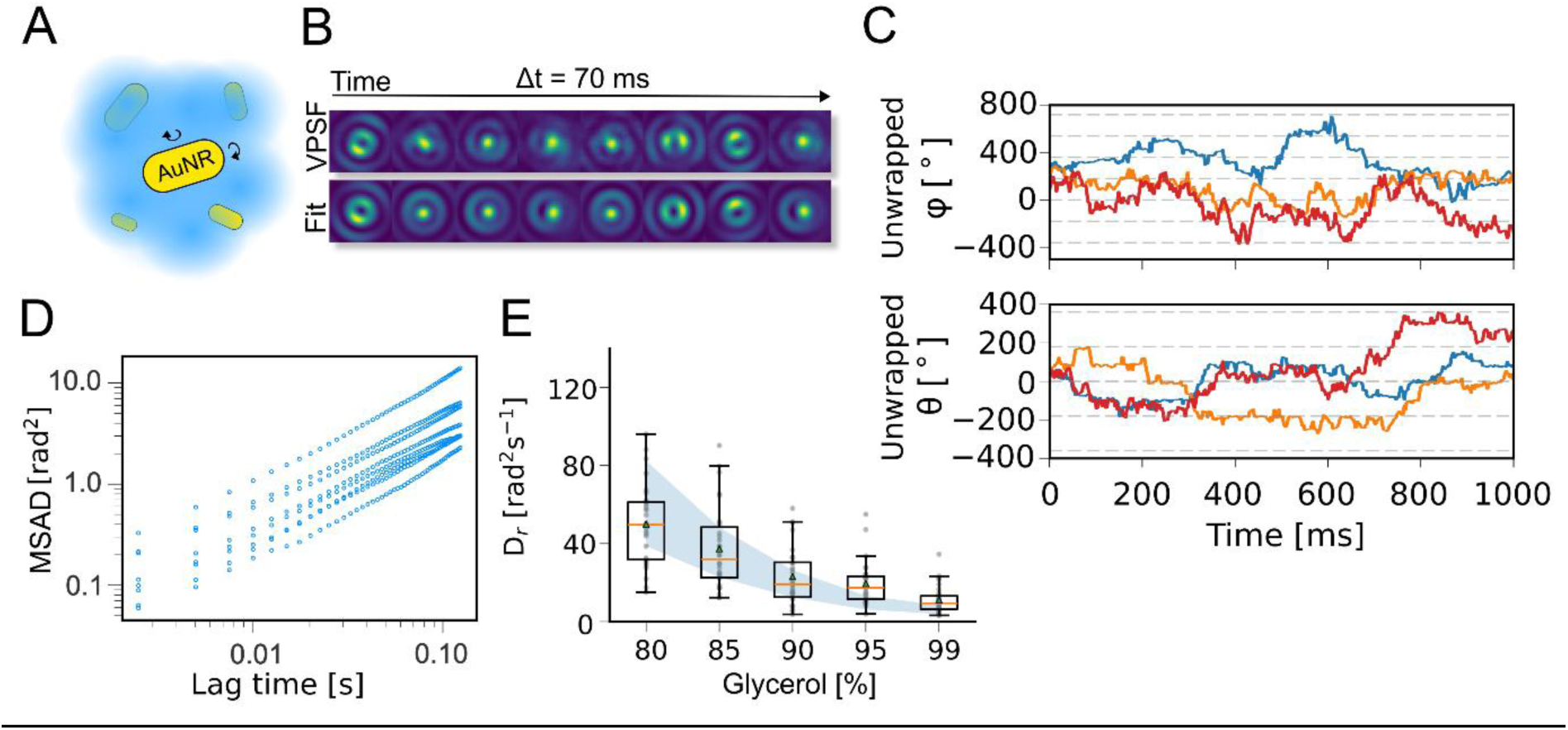
| Quantitative measurement of rotational diffusion using freely diffusing AuNRs. **(A)** Schematic of freely diffusing AuNRs in glycerol–water mixtures. **(B)** Time-lapse montage of VPSFs and corresponding orientation fits for a representative freely diffusing AuNR in an 80% glycerol–water mixture. **(C)** Representative unwrapped azimuthal (ϕ) and polar (θ) trajectories for three freely diffusing AuNRs in an 80% glycerol–water mixture. **(D)** Representative mean squared angular displacement (MSAD) curves for freely diffusing AuNRs in 99% glycerol. **(E)** Rotational diffusion coefficients measured as a function of glycerol concentration (80–99% glycerol). Blue shading indicates the theoretical prediction ± the uncertainty arising from the AuNR size distribution.

For freely diffusing particles, the MSAD increases linearly with lag time (4*D_r_*Δ*_t_*), allowing direct determination of D_ᵣ_. Across the viscosity range examined, the measured rotational diffusion coefficients closely matched theoretical predictions for rod-like particles (Fig. 2E; Methods) [^24,25^], demonstrating accurate quantitative measurements of rotational diffusion.

### Translational and rotational diffusion provide complementary information about the local physical environment

Having established accurate measurements of rotational diffusion, we next asked whether simultaneous measurements of translational and rotational diffusion provide complementary information about the local physical environment. To address this question, we tethered AuNRs to supported lipid bilayers via biotin–streptavidin linkages and monitored their translational and rotational dynamics (Fig. 3A-C; Fig. S3C,D, Supplementary Video S1).

**Figure 3.**
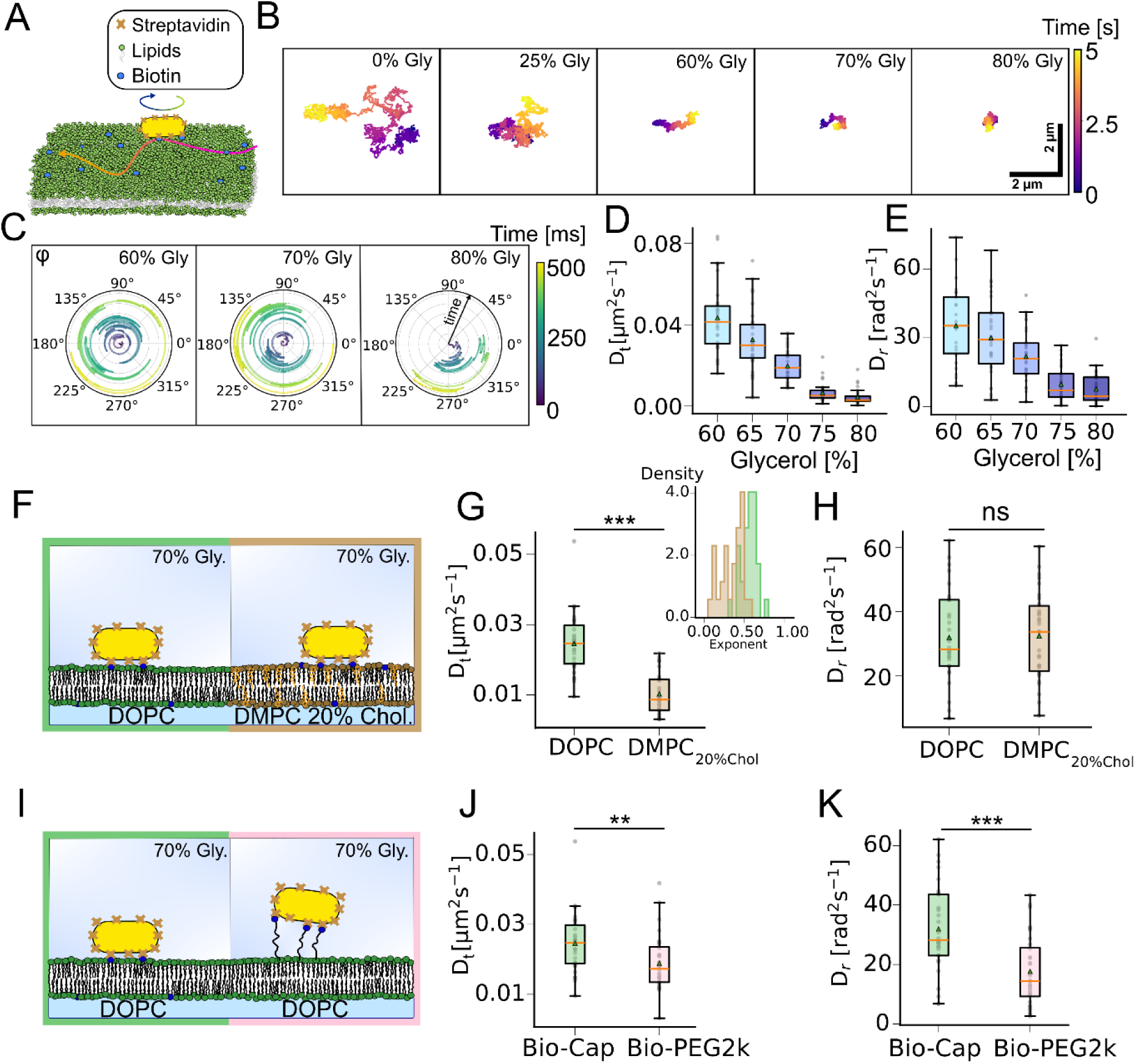
|Translational and rotational diffusion provide complementary readouts of the local physical environment. **(A)** Schematic of an AuNR tethered to a supported lipid bilayer via biotinylated lipids. **(B)** Representative translational trajectories of single AuNRs on a DOPC bilayer measured at increasing glycerol concentrations. **(C)** Representative polar plots of the azimuthal angle (ϕ) for single AuNRs on a DOPC bilayer measured at increasing glycerol concentrations. The radial coordinate represents time over a total trajectory duration of 500 ms. **(D,E)** Translational (D) and Rotational (E) diffusion coefficients of AuNRs on a DOPC bilayer measured across 60–80% glycerol-water mixtures. **(F)** Schematic of AuNRs tethered to DOPC or DMPC + 20% cholesterol bilayers in 70% glycerol. **(G,H)** Translational (G) and rotational (H) diffusion coefficients of AuNRs on DOPC and DMPC + 20% cholesterol bilayers. Panel (G) additionally features the MSD exponent histograms. **(I)** Schematic of AuNRs tethered to a DOPC bilayer via short biotinyl-cap-PE or long DSPE-PEG2000-biotin linkers in 70% glycerol. **(J,K)** Translational (J) and rotational (K) diffusion coefficients of AuNRs tethered via biotinyl-cap-PE or DSPE-PEG2000-biotin. P values in panels G, H, J, and K were calculated using two-sided *t*-tests.

Increasing glycerol concentration progressively reduced both translational and rotational diffusion coefficients (Fig. 3D,E), consistent with the increase in bulk viscosity. To verify accurate measurements of rotational diffusion over the accessible dynamic range, we compared rotational diffusion coefficients obtained from MSAD analysis with those derived from frame-to-frame angular displacement distributions (Fig. S3E-G). Above approximately 60% glycerol, both approaches converged, confirming accurate quantification of rotational diffusion.

We next altered membrane fluidity by comparing fluid DOPC bilayers with DMPC bilayers containing 20% cholesterol, which exhibit increased lipid order and reduced membrane fluidity (Fig. 3F). As expected, translational diffusion was significantly reduced in the more ordered membrane (Fig. 3G), consistent with independent FRAP measurements (Fig. S4A–F). Mean-squared displacement analysis further revealed stronger sub-diffusive behavior, reflecting reduced membrane mobility (Fig. 3G). In contrast, rotational diffusion coefficients remained essentially unchanged (Fig. 3H).

Finally, we examined how nanoparticle–membrane coupling influences translational and rotational dynamics by increasing the linker length between the nanoparticle and the membrane (Fig.3I). Increasing the linker length reduced both translational and rotational diffusion coefficients (Fig. 3J,K), with a substantially larger effect on rotational diffusion, consistent with stronger coupling between the nanoparticle and its local environment.

Together, these experiments establish translational and rotational diffusion as complementary reporters of the local physical environment. Whereas translational diffusion reflects both bulk viscosity and membrane mobility, rotational diffusion is largely insensitive to membrane mobility and instead reports the immediate local environment surrounding the nanoparticle. Furthermore, increasing the linker length enhances the sensitivity of rotational diffusion to this local environment. Combining these complementary observables therefore provides biophysical information that cannot be obtained from translational tracking alone.

### Simultaneous translational and rotational tracking distinguishes intracellular environments

We next asked whether these complementary measurements could distinguish the endolysosomal system from the cytosol, two intracellular environments with distinct physical and biochemical properties [^26–28]^. To address this, AuNRs were introduced into living HeLa cells either directly into the cytoplasm by transient surfactant-mediated permeabilization or through endocytic uptake into the endolysosomal system (Fig. 4A, Supplementary Video S2, Methods), and their translational and rotational dynamics were compared. Endolysosomal compartments were labeled with LysoBrite Red. Cytoplasmically delivered AuNRs showed minimal colocalization with LysoBrite-positive structures (6%, N=2 out of 32), whereas endocytosed AuNRs showed complete colocalization with LysoBrite-labeled compartments (100%, N=69), confirming their distinct intracellular localization (Fig. 4B,C). In situ cryo-electron tomography (cryo-ET) independently confirmed that endocytosed AuNRs were enclosed within endolysosomal compartments (Fig. 4D, S5A-F)[^29^]. Cryo-ET was performed at a substantially higher AuNR concentration than single-particle tracking to increase the likelihood of capturing nanorods within the tomographic volume. As a result, multiple nanorods were occasionally observed within the same compartment.

**Figure 4.**
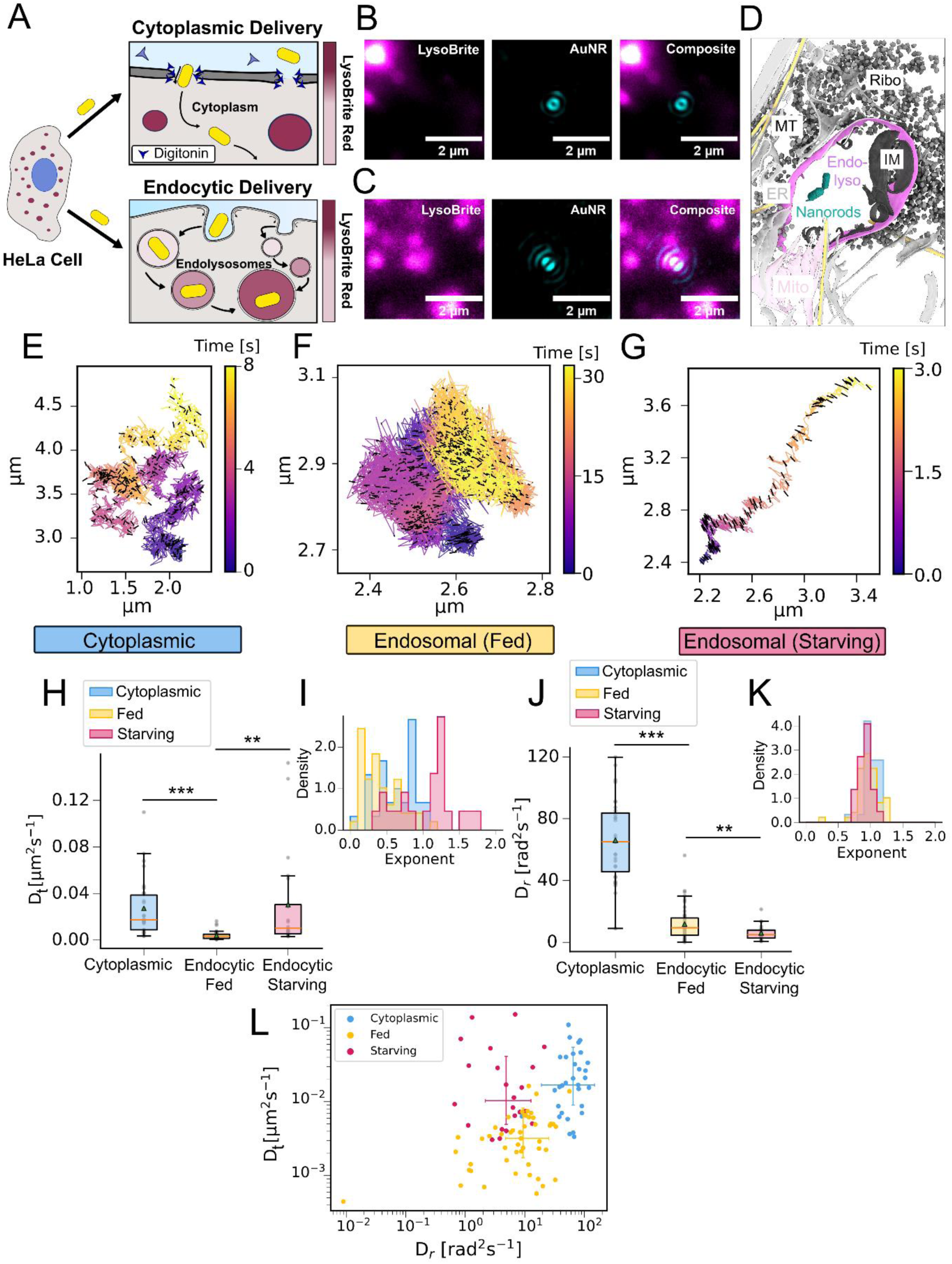
| Multidimensional single-particle tracking resolves intracellular environments and physiological remodeling. **(A)** Schematic of cytoplasmic delivery of AuNRs by digitonin permeabilization (top) and endocytic uptake (bottom) in HeLa cells. **(B,C)** Representative fluorescence images of LysoBrite-labeled compartments and the corresponding scattering images of a cytoplasmically delivered (B) and an endocytosed (C) AuNRs. Merged images show the absence (B) or presence (C) of colocalization. **(D)** Segmented cryo-electron tomography reconstruction showing AuNRs enclosed within an endo-lysosomal compartment following endocytic uptake. **(E–G)** Representative trajectories of cytoplasmically delivered AuNRs (E), endocytosed AuNRs in nutrient-fed cells (F), and endocytosed AuNRs in starved cells (G). Tick orientation indicates the azimuthal angle (ϕ), and tick length represents the projected polar angle, 1 − sin(θ). **(H,J)** Boxplots of translational (H) and rotational (I) diffusion coefficients for cytoplasmically delivered AuNRs, endocytosed AuNRs in nutrient-fed cells, and endocytosed AuNRs in starved cells. **(I,K)** Histograms of anomalous diffusion exponents derived from translational mean squared displacement (MSD; I) and mean squared angular displacement (MSAD; K) analyses for the three conditions. **(L)** Scatter plot of rotational (Dr) versus translational (Dt) diffusion coefficients showing three distinct intracellular populations. Error bars indicate the interquartile ranges centered on the median values. Statistical significance was assessed using two-sided *t*-tests.

Representative trajectories (Fig. 4E,F) together with quantitative analysis of translational (D_t_) and rotational (D_r_) diffusion coefficients (Fig. 4H,J) revealed that endocytosed AuNRs exhibited significantly slower translational and rotational motion than cytoplasmic particles, consistent with the distinct physical properties of the membrane-enclosed endolysosome [^30^]. Mean-squared displacement analysis showed predominantly subdiffusive translational motion (exponent < 1) in both the cytoplasm and endolysosomes, with stronger subdiffusion in endolysosomes, indicating increased confinement and restricted translational mobility within membrane-bound compartments (Fig. 4I).

In contrast, the rotational diffusion exponent remained close to one under both conditions (Fig. 4K), indicating that rotational motion remained Brownian despite the substantial reduction in rotational diffusivity. The decrease in D_r_ despite a Brownian rotational exponent suggests that the slower rotational diffusion reflects an increase in local microviscosity or frictional drag, rather than geometric confinement or transient trapping of the nanorod. This observation is consistent with the cryo-ET data showing that AuNRs remained freely suspended within endolysosomal compartments rather than attached to the membrane (Fig. 4D). Together, these results demonstrate that simultaneous measurements of D_t_ and D_r_ clearly distinguish cytoplasmic and endolysosomal nanoparticles in two-dimensional diffusion space (Fig. 4L), demonstrating that translational and rotational diffusion provide complementary information for characterizing intracellular microenvironments.

We next asked whether these complementary measurements could also detect physiological remodeling of intracellular environments. Nutrient deprivation is known to remodel the endolysosomal system as it stimulates endolysosomal movement towards the perinuclear region of the cell [^31–34]^. To test this we compared AuNR-containing endocytic compartments in cells maintained in complete medium with cells starved for 1–2 h in PBS (Fig. 4F,G)

Because translational motion of endocytosed nanoparticles reflects both compartment transport and nanoparticle motion within the compartment, we first independently assessed endolyososmal movement by tracking LysoBrite-positive endolysosomal compartments. In line with previous observations, extended starvation significantly increased endolysosomal mobility (Fig. S5G). Consistent with this observation, endocytosed AuNRs exhibited significantly higher translational diffusion coefficients under starvation (Fig. 4G,H). Representative trajectories frequently displayed directional movement (Fig.4G), and a substantial fraction of endolysosomes under extended starvation condition exhibited superdiffusive behavior (Exponent > 1, Fig.4I), consistent with active intracellular transport [^35^].

In contrast, rotational diffusion decreased significantly under the same conditions (Fig. 4J), despite the increased mobility of the compartments. Notably, the rotational diffusion exponent remained close to one (Fig. 4K), suggesting a further increase in microviscosity. This is in line with a previous study that during starvation lysosome viscosity increases [^36^]. These opposing changes demonstrate that translational and rotational diffusion report complementary aspects of endolysosomal dynamics: translational diffusion primarily reflects compartment transport, whereas rotational diffusion reports the local nanoscale physical environment. Consequently, fed and extended-starved endolysosome occupied distinct regions in the two-dimensional diffusion space (Fig. 4L), highlighting the potential of combined diffusion measurements for biophysical fingerprinting physiological remodeling of intracellular microenvironments.

### Temporal remodeling within individual endosomes

Finally, we asked whether these complementary measurements could resolve temporal remodeling within individual endolysosomes. To capture such dynamic changes, individual trajectories were divided into consecutive time windows, from which translational and rotational diffusion coefficients were calculated independently (Fig. 5A-C). Representative trajectories of endocytosed AuNRs in starved cells revealed pronounced fluctuations in both translational and rotational motions during the tracking period (Fig. 5B-D). Notably, changes in translational mobility were frequently not accompanied by corresponding changes in rotational diffusivity, and vice versa, indicating that the two observables report distinct dynamic processes within individual endocytic compartments.

**Figure 5.**
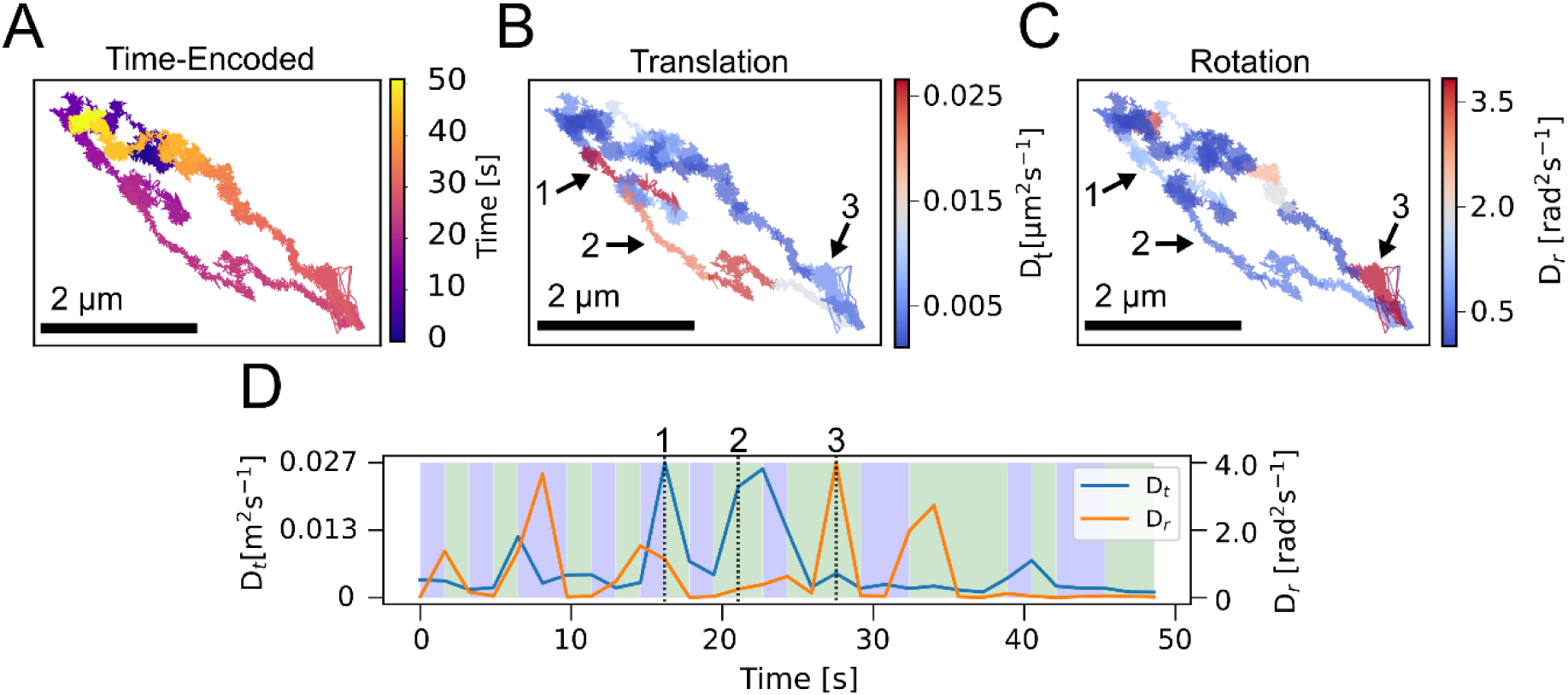
| Time-resolved multidimensional single-particle tracking reveals dynamic remodeling of the local intracellular environment. **(A)** Time-encoded trajectory of a single endocytosed AuNR in a starved HeLa cell. **(B,C)** Time evolution of translational (B) and rotational (C) diffusion along the trajectory shown in (A). **(D)** Time traces of the translational (Dt, blue) and rotational (Dr, orange) diffusion coefficients. Green shaded regions indicate coordinated changes in both coefficients, whereas purple shaded regions indicate opposing changes over time.

Together, these results establish combined translational and rotational measurements as a framework for probing intracellular environments across multiple scales. By combining translational and rotational diffusion measurements, the approach distinguishes intracellular environments, detects physiological remodeling, and resolves dynamic changes within individual compartments that are inaccessible to conventional translational tracking.

## Discussion

Here, we introduce a tracking approach that simultaneously measures the translational and rotational dynamics of individual nanoparticles to probe intracellular environments. By combining these complementary observables, the approach distinguishes intracellular environments, uncovers starvation-induced remodeling of endocytic compartments, and resolves dynamic changes within individual compartments over time that are inaccessible through conventional positional tracking alone.

Experiments in a controlled membrane system demonstrated that translational and rotational diffusion respond differently to distinct physical perturbations. Whereas translational diffusion primarily reflects the motion of the structure to which the nanoparticle is coupled, rotational diffusion is more strongly influenced by the nanoparticle’s immediate nanoscale surroundings. These distinct sensitivities establish translational and rotational dynamics as complementary observables that report different physical aspects of the local environment.

Applying this approach to living cells demonstrated that translational and rotational dynamics together distinguish distinct intracellular environments. Nanoparticles delivered directly into the cytoplasm exhibited substantially faster translational and rotational dynamics than particles confined within endocytic compartments, confirming that the combined measurements readily distinguish distinct intracellular physical environments. Within the endocytic pathway itself, nutrient starvation further revealed a striking decoupling between compartment transport and the local environment, with increased compartment mobility accompanied by reduced rotational diffusion. Time-resolved analysis further showed that changes in translational mobility frequently occurred independently of changes in rotational diffusion within individual endocytic compartments, indicating that local environmental remodeling can proceed independently of compartment transport. Although the molecular basis of these observations remains to be determined, they could arise from changes in molecular crowding, local viscosity, steric confinement, or nanoparticle–compartment interactions.

Together, these findings demonstrate that combining translational and rotational tracking resolves both large-scale intracellular transport and local nanoscale physical changes within the same experiment. More broadly, this approach provides a versatile approach for independently interrogating the dynamics of biological structures and the physical properties of their local environment. A key advantage of this framework is that the balance between sensitivity to structural dynamics and the local physical environment can be tuned through the design of the nanoparticle– target interface. For example, increasing linker length enhances sensitivity to the surrounding medium, whereas increasing the number of attachment points progressively restricts rotational freedom (Fig. S3H-K), allowing rotational motion to more faithfully report the dynamics of the underlying structure. This design flexibility should facilitate adaptation of the approach to a wide range of biological systems, including membranes, intracellular organelles, cytoskeletal assemblies, biomolecular condensates, and chromatin.

The widespread adoption of this approach will depend on continued advances in several complementary areas. Reliable nanoparticle delivery, intracellular targeting strategies, and biocompatible probe design will be essential for extending the approach to specific biological structures while preserving their native dynamics. Equally important will be the implementation of multidimensional tracking on widely available microscopy platforms, together with fast, automated analysis pipelines capable of routine extraction of translational and rotational dynamics from large datasets. Future developments, including three-dimensional localization of AuNRs and experimental and analytical methods to extract sub-frame rotational dynamics [^37^] could further expand the versatility of the approach. As these technological advances continue, this approach should become increasingly accessible across the biological sciences. By simultaneously resolving the dynamics of biological structures and the physical properties surrounding them, the approach offers a new lens for uncovering how physical organization shapes cellular behavior across spatial and temporal scales.

## Data and Materials Availability

All data are available upon request to the corresponding authors. The analysis code will be available in the following link upon publication: https://github.com/DimitriSchumacher

## Acknowledgements

D.S was supported by the Deutsche Forschungsgemeinschaft (DFG, German Research Foundation)-SPP2202. Scientific support was provided by Kestrel Neuroscience (W.Z.). D.L. was supported by a Boehringer Ingelheim Fonds PhD Fellowship. The work was supported by the Max Planck Society, the European Union (F.W.: ERC, IntrinsicReceptors, 101041982, E.K: ERCStG LoopSMC 101076914). Views and opinions expressed are, however, those of the author(s) only and do not necessarily reflect those of the European Union or the ERC Executive Agency. Neither the European Union nor the granting authority can be held responsible for them.

## Author Contributions

M.D.B. conceived the initial idea for the methodology. D.S., M.D.B., E. K. developed the methodology. W.Z. performed cytoplasmic and endocytic NR deliveries and did cell culture and cell preparations. D.L. performed cryo-ET measurements. D.S., M.D.B., W.Z., B.P., D.L., F.W., and E.K. performed the investigation. D.S., M.D.B., W.Z., D.L., and T.F. contributed to data visualization. F.W. and E.K. acquired funding and administered the project. D.S., E.K. wrote the initial draft of the manuscript. D.S., M.D.B., W.Z., B.P., D.L., F.W., and E.K. contributed to reviewing and editing the manuscript.

## Methods

### Microscope design

The microscope was built in a widefield illumination configuration using 561 nm and 638 nm lasers (Cobolt o6-DPL-561 and Cobolt o6-MLD-638). The collimated excitation beam was focused at the back focal plane of the objective to generate widefield illumination, and circular polarization was achieved using a quarter-wave plate. A 4f relay system (f = 165 mm) was incorporated into the emission path to position a vortex phase plate (Thorlabs, m = 1 vortex retarder, optimized for 633 nm) at a plane conjugate to the objective back focal plane (Zeiss Alpha Plan-Apochromat 100×/1.46 Oil). A custom 5 mm diameter beam stop was introduced in the emission path to create a near-darkfield imaging configuration for suppression of background scattering. A schematic of the optical setup is provided in Supplementary Fig. S1.

### PSF simulation

Point spread functions (PSFs) were simulated using vectorial diffraction theory as previously described [^38^]. A detailed description of the simulation framework is provided in the Supplementary Information.

### Brute-force fitting routine

We developed a GPU-accelerated brute-force fitting routine in Python using the CuPy library [^39^]. The algorithm compares each experimentally acquired vortex point spread function (VPSF) with a precomputed library of simulated VPSFs spanning all combinations of the azimuthal (ϕ) and polar (θ) angles at 1° angular increments. For each orientation, the least-squares residual between the experimental and simulated VPSFs is calculated. To account for lateral misalignment between the experimental VPSF and the simulated library, image registration is performed by evaluating the residuals over a defined range of x–y offsets. The orientation and registration parameters corresponding to the global minimum residual are selected as the optimal fit. The goodness of fit for each frame is quantified using the coefficient of determination (R²). Further details of the fitting algorithm are provided in the Supplementary Information.

### Analysis pipeline

The analysis pipeline, from raw image acquisition to translational and rotational diffusion analysis, is described in detail in the Supplementary Information. Briefly, vortex point spread functions (VPSFs) were first extracted and centered from the raw image sequences using a custom Python program implemented in Napari [^40^]. VPSFs were localized using a series of image-processing steps, including inverted Sato filtering (Scikit-Image [^41^]) and intensity thresholding, and cropped into 31 × 31 pixel regions for subsequent analysis. Individual VPSFs were then fitted using the GPU-accelerated fitting algorithm described above to determine their orientations, while the corresponding centroids were used for translational tracking. The resulting angular trajectories were unwrapped using Numpy’s unwrap function with periods of 2π for the azimuthal angle (ϕ) and π for the polar angle (θ) [^42^]. Mean squared displacement (MSD) and mean squared angular displacement (MSAD) analyses were subsequently performed using the centroid and unwrapped angular trajectories, respectively. Diffusive behavior was quantified by fitting the MSD and MSAD curves with the power-law model, *y* = *Kx***^α^**. Further details of the analysis pipeline are provided in the Supplementary Information.

### Nanostructured AuNRs

Gold nanorods were fabricated by focused ion beam (FIB) milling of wet-chemically grown 50 nm-thick gold flakes. Gold flakes were transferred onto glass coverslips (Menzel, 22 × 22 mm, No. 1.5) using a poly(methyl methacrylate) (PMMA)-assisted transfer method. The coverslips were pre-patterned with evaporated gold hole masks for localization. Nanorod outlines were milled using a Ga-ion FIB microscope (FEI Helios NanoLab), after which the remaining gold film was removed by adhesive tape lift-off, leaving only the nanostructured AuNRs on the substrate.

### Luminescence imaging

Luminescence images of AuNRs on prefabricated nanogrids were acquired using a 561 nm laser (Cobolt o6-DPL-561) operated at 120 mW with an exposure time of 2 s per frame.

### Fluorescence imaging

Fluorescence images of supported lipid bilayers and LysoBrite-labeled HeLa cell compartments were acquired using a 561 nm laser (Cobolt o6-DPL-561) operated at 25 mW with an exposure time of 100 ms.

### Scattering imaging

AuNR scattering images were acquired using a 638 nm laser (Cobolt o6-MLD-638) operated at 180 mW. Images were recorded with a 1 ms camera exposure to maximize the acquisition rate. For experiments involving translational motion, a 256 × 256 pixel region of interest (ROI) was used, resulting in an average frame interval of approximately 2.5 ms.

### Coverslip cleaning

Unfunctionalized glass coverslips were cleaned by alternately rinsing with Milli-Q water and isopropanol five times. During each rinse, the solvent was applied from the top of the coverslip and allowed to flow across the surface to ensure complete coverage. After the final rinse, the coverslips were dried under a gentle nitrogen stream and immediately used for flow-cell assembly.

### Surface immobilization of AuNRs

To obtain a homogeneous distribution of surface-immobilized AuNRs, cleaned glass flow cells were first rinsed with 100 μL Milli-Q water. CTAB-coated AuNRs (Nanopartz, A12-40-650-CTAB-DIH-1-25) were diluted 1:10 in Milli-Q water (final concentration approximately 120 pM) to reduce the residual CTAB concentration, which otherwise inhibits surface adsorption. The diluted AuNRs were introduced into the flow cell and incubated for up to 5 min to allow adsorption to the glass surface. Immobilization was then induced by flushing the channel with 100 μL of 5 M NaCl.

### Theoretical rotational diffusion

Theoretical calculations of rotational diffusion coefficients for freely diffusing AuNRs were made with the following equation [^24^],

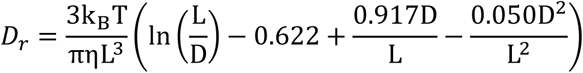

where k_B_ is the Boltzmann constant, η is the viscosity, T is the temperature, L is the length of the AuNR and D its diameter.

### Supported lipid bilayers

Supported lipid bilayers were prepared using the solvent-assisted lipid bilayer (SALB) method [^43,44^]. Lipid mixtures consisted of DOPC or DMPC supplemented with biotinyl-cap-PE, DSPE-PEG2000-biotin, and 0.25 mol% Liss-Rhod-PE for fluorescence imaging (Avanti-Research). Where indicated, 20 mol% cholesterol was included. Lipids were mixed in chloroform, dried under nitrogen, and redissolved in isopropanol (2.5 mg mL⁻¹) immediately before bilayer formation.

Bilayers were formed in cleaned glass flow cells using a syringe pump at a flow rate of 50 μL min⁻¹ by sequential exchange of Tris-NaCl buffer (10 mM Tris, 150 mM NaCl, pH 7.5), isopropanol, lipid solution, and buffer. After a 5 min incubation with the lipid solution, excess lipid was removed by buffer exchange.

For AuNR tethering, streptavidin-functionalized biotin-AuNRs (Nanopartz, C12-40-650-TB-DIH-50-1) were diluted to 400 pM in PBS, introduced into the flow cell, and incubated until the desired particle density was reached. Unbound particles were removed by washing with PBS. Where indicated, PBS containing glycerol was introduced at 15 μL min⁻^¹^ to achieve the desired glycerol concentration.

### Cell culture and live-cell imaging

HeLa cells (CCL-2; RRID:CVCL_0030) were maintained in Dulbecco’s modified Eagle medium supplemented with 10% fetal bovine serum, 4 mM L-glutamine, and 1% penicillin–streptomycin at 37 °C in a humidified incubator with 5% CO_₂_. Cells were seeded at 40,000 cells per well in glass-bottom 8-well μ-Slides (ibidi, 80807) and cultured overnight before imaging.

For endocytic uptake experiments, cells were incubated with 5–10 pM carboxylated AuNRs (Nanopartz, C12-40-650-TC-DIH-50-1) for 4 h before imaging. Cytoplasmic delivery of AuNRs was performed using a modified digitonin permeabilization protocol as described previously [^45^]. Briefly, cells were permeabilized with 10 μg mL⁻¹ digitonin in DPBS for 10 min at 4 °C, washed with DPBS supplemented with 0.9 mM CaCl₂ and 0.49 mM MgCl₂, and incubated with 5 pM AuNRs for 30 min at 37 °C. Cells were then allowed to recover for 1 h in complete medium without penicillin–streptomycin.

Lysosomes were labeled with LysoBrite Red (1:2500; AAT Bioquest, 22645) for 10 min before imaging. Live-cell imaging was performed in DPBS supplemented with 15% iodixanol to improve optical image quality [^46^].

### *In situ* cryogenic electron tomography (cryo-ET) of AuNRs in cellular compartments

#### Sample preparation

HeLa cells with endocytosed AuNRs (Nanopartz, C12-40-650-TC-DIH-50-1) were seeded for *in situ* cryo-ET as follows: Plasma-cleaned (Pelco easiglow, 2×90 s, 15 mA, 0.37 mbar) Quantifoil SiO_2_/Au R1/4 mesh 200 EM grids were coated with 0.25 µg/mL laminin (Sigma-Aldrich, L4544) for 1 h and UV sterilized. The grids were washed twice in PBS before transferring into 8-well µ-Slide dishes (Ibidi, 80826). 10,000 cells were seeded on top of the grids in DMEM high glucose HEPES medium without phenol red (Thermo Fisher Scientific, 21063029) and incubated overnight at 37 °C and 5% CO_2_. The next day, the medium was replaced with medium supplemented with 800 pM AuNR for 4 h then washed with complete medium and vitrified using a Leica EM GP2 (Leica Microsystems) plunge freezer (blot time 4 s, with the chamber set to 37 °C and 70 % humidity) in liquid ethane at -184 °C. 5 min before plunge-freezing LysoBrite Red (1:4000) was added. Vitrified grids were clipped into autogrids with a cutoff for FIB-milling.

#### Focused ion beam (FIB) milling

150 nm to 200 nm thick lamella of HeLa cells were produced in a Aquilos2 (Thermo Fisher Scientific) dual-beam cryo-focused ion beam and scanning electron microscope (cryo-FIB/SEM). Grids were coated with a layer of organometallic platinum using the built-in gas injection system (GIS) for 45 s and afterwards an inorganic platinum layer was applied using the sputter coater for 20 s. Grid maps were acquired using MAPS (v.3.28, Thermo Fisher Scientific). Semi-automated FIB-milling was done using AutoTEM (v.2.4, Thermo Fisher Scientific) in the following steps: i) rough milling at 1 nA to 1 µm thickness, (ii) 0.5 nA to 750 nm, and (iii) 0.3 nA to the 500 nm. Afterwards, the lamella was polished at a current of 50 pA to 150 nm and 10 pA to 130 nm (with 0.2° overtilting). Manual polishing was carried out to further thin down lamella and thickness was evaluated using the SEM at 3 kV, 13 pA.

#### Tilt series acquisition

Tilt series of AuNR in cellular compartments were acquired on a Titan Krios G2 (Thermo Fisher Scientific) cryogenic transmission electron microscope (cryo-TEM) operated at 300 kV equipped with a BioQuantum energy filter set to a slit width of 20 eV and a K3 direct electron detector (Gatan) at a nominal magnification of 33,000 X corresponding to a pixel size of 2.682 Å/px at bin1. The acquisition was controlled using SerialEM (v.4.1) [^47^] with the target defocus ranging from -3 µm to -5 µm and the dose set to ∼150 e/Å^2^ spread over 60 tilts ranging from +52° to -66° to compensate for the lamella pre-tilt (-8°). 10 frames per individual tilt were acquired and on-the-fly motion corrected in SerialEM. The positions for tilt series acquisition were determined based on the lamella maps acquired at 8,700 X.

#### Tomogram reconstruction and denoising

The tilt series mrc stacks were corrected for dose exposure and projections of low-quality were removed after manual inspection. The tilt series were automatically aligned using patch-tracking and reconstructed by weighted back-projection at bin4 using AreTomo2 (v.1.0.0) [^48^] and IMOD (v.4.12.65) [^49^].

For denoising, tomograms were reconstructed in similar fashion starting from stacks that contained either odd or even frames. The split tomograms were reconstructed and subjected to denoising with Isonet2 [^50^] using default parameters.

All membranes in the tomograms were segmented using Membrain-Seg v.0.0.8 [^51^] using the pre-trained model (v10_alpha). Segmentations were touched up and cleaned manually in Napari [^40^] to correct falsely connected membranes. All visualization renders were done using ChimeraX v.1.11. The nanorods were segmented by just intensity thresholding the darkest 99.9% of pixels in the tomogram. Ribosomes and microtubules were segmented using easymode [^52^].

## References

1. Simon, F., Weiss, L. E. & van Teeffelen, S. A guide to single-particle tracking. Nat. Rev. Methods Primer 4, 66 (2024).

2. Scheiderer, L., Marin, Z. & Ries, J. MINFLUX achieves molecular resolution with minimal photons. Nat. Photonics 19, 238–247 (2025).

3. Tirado, M. M., Martínez, C. L. & De La Torre, J. G. Comparison of theories for the translational and rotational diffusion coefficients of rod-like macromolecules. Application to short DNA fragments. J. Chem. Phys. 81, 2047–2052 (1984).

4. Garcia de la Torre, J. G. & Bloomfield, V. A. Hydrodynamic properties of complex, rigid, biological macromolecules: theory and applications. Q. Rev. Biophys. 14, 81–139 (1981).

5. Gu, Y. et al. Single Particle Orientation and Rotational Tracking (SPORT) in biophysical studies. Nanoscale 5, 10753–10764 (2013).

6. Anthony, S. M. & Yu, Y. Tracking single particle rotation: probing dynamics in four dimensions. Anal. Methods 7, 7020–7028 (2015).

7. Senthil Kumar, C. S., et al. 4polar3D single molecule imaging of 3D orientation in dense actin networks using ratiometric polarization splitting. Nat. Commun. https://doi.org/10.1038/s41467-026-70852-y (2026) doi:10.1038/s41467-026-70852-y.

8. Sun, B., Ding, T., Zhou, W., Porter, T. S. & Lew, M. D. Single-Molecule Orientation Imaging Reveals the Nano-Architecture of Amyloid Fibrils Undergoing Growth and Decay. Nano Lett. 24, 7276–7283 (2024).

9. Sarkar, A., Mitra, J. B., Sharma, V. K., Namboodiri, V. & Kumbhakar, M. Spectrally Resolved Single-Molecule Orientation Imaging Reveals a Direct Correspondence between the Polarity and Microviscosity Experienced by Nile Red in Supported Lipid Bilayer Membranes. J. Phys. Chem. B 129, 2380–2391 (2025).

10. Bruggeman, E. et al. POLCAM: instant molecular orientation microscopy for the life sciences. Nat. Methods 21, 1873–1883 (2024).

11. Brasselet, S. & Lew, M. D. Single-molecule orientation and localization microscopy. Nat. Photonics 19, 925–937 (2025).

12. Zhang, O. et al. Six-dimensional single-molecule imaging with isotropic resolution using a multi-view reflector microscope. Nat. Photonics 1–8 (2022) doi:10.1038/s41566-022-01116-6.

13. Hulleman, C. N. et al. Simultaneous orientation and 3D localization microscopy with a Vortex point spread function. Nat. Commun. 12, 5934 (2021).

14. Chaudhari, K. & Pradeep, T. Spatiotemporal mapping of three dimensional rotational dynamics of single ultrasmall gold nanorods. Sci. Rep. 4, 5948 (2014).

15. Chen, J. & Irudayaraj, J. Quantitative Investigation of Compartmentalized Dynamics of ErbB2 Targeting Gold Nanorods in Live Cells by Single Molecule Spectroscopy. ACS Nano 3, 4071–4079 (2009).

16. Austin, L. A., Kang, B. & El-Sayed, M. A. Probing molecular cell event dynamics at the single-cell level with targeted plasmonic gold nanoparticles: A review. Nano Today 10, 542–558 (2015).

17. Ye, W. et al. Conformational Dynamics of a Single Protein Monitored for 24 h at Video Rate. Nano Lett. 18, 6633–6637 (2018).

18. Mazaheri, M., Ehrig, J., Shkarin, A., Zaburdaev, V. & Sandoghdar, V. Ultrahigh-Speed Imaging of Rotational Diffusion on a Lipid Bilayer. Nano Lett. 20, 7213–7219 (2020).

19. Sönnichsen, C. & Alivisatos, A. P. Gold Nanorods as Novel Nonbleaching Plasmon-Based Orientation Sensors for Polarized Single-Particle Microscopy. Nano Lett. 5, 301–304 (2005).

20. Cole, D., Young, G., Weigel, A., Sebesta, A. & Kukura, P. Label-Free Single-Molecule Imaging with Numerical-Aperture-Shaped Interferometric Scattering Microscopy. ACS Photonics 4, 211–216 (2017).

21. Liebel, M., Hugall, J. T. & van Hulst, N. F. Ultrasensitive Label-Free Nanosensing and High-Speed Tracking of Single Proteins. Nano Lett. 17, 1277–1281 (2017).

22. Einstein, A. Über die von der molekularkinetischen Theorie der Wärme geforderte Bewegung von in ruhenden Flüssigkeiten suspendierten Teilchen. Ann. Phys. 322, 549–560 (1905).

23. Qian, H., Sheetz, M. P. & Elson, E. L. Single particle tracking. Analysis of diffusion and flow in two-dimensional systems. Biophys. J. 60, 910–921 (1991).

24. Tirado, M. M. & De La Torre, J. G. Rotational dynamics of rigid, symmetric top macromolecules. Application to circular cylinders. J. Chem. Phys. 73, 1986–1993 (1980).

25. Volk, A. & Kähler, C. J. Density model for aqueous glycerol solutions. Exp. Fluids 59, 75 (2018).

26. Luby-Phelps, K. Cytoarchitecture and physical properties of cytoplasm: volume, viscosity, diffusion, intracellular surface area. Int. Rev. Cytol. 192, 189–221 (2000).

27. Kalwarczyk, T. et al. Comparative Analysis of Viscosity of Complex Liquids and Cytoplasm of Mammalian Cells at the Nanoscale. Nano Lett. 11, 2157–2163 (2011).

28. Chambers, J. E. et al. An Optical Technique for Mapping Microviscosity Dynamics in Cellular Organelles. ACS Nano 12, 4398–4407 (2018).

29. Li, D. et al. Cathepsin-dependent amyloid formation drives mechanical rupture of lysosomal membranes. bioRxiv 2026.01.17.700056 (2026) doi:10.64898/2026.01.17.700056.

30. Chen, K. et al. Characteristic rotational behaviors of rod-shaped cargo revealed by automated five-dimensional single particle tracking. Nat. Commun. 8, 887 (2017).

31. Korolchuk, V. I. et al. Lysosomal positioning coordinates cellular nutrient responses. Nat. Cell Biol. 13, 453–460 (2011).

32. Ebner, M. et al. Nutrient-regulated control of lysosome function by signaling lipid conversion. Cell 186, 5328–5346.e26 (2023).

33. Sareen, S., Zgorzelska, A., Kwapiszewska, K. & Hołyst, R. Starvation induces diffusion hindrance at the nanoscale in mammalian cells. Nanoscale 17, 378–389 (2024).

34. Shi, H. et al. Starvation induces shrinkage of the bacterial cytoplasm. Proc. Natl. Acad. Sci. U. S. A. 118, e2104686118 (2021).

35. Guo, Y. et al. Visualizing Intracellular Organelle and Cytoskeletal Interactions at Nanoscale Resolution on Millisecond Timescales. Cell 175, 1430–1442.e17 (2018).

36. Li, D. et al. Nucleic acid-selective light-up fluorescent biosensors for ratiometric two-photon imaging of the viscosity of live cells and tissues. Chem. Sci. 7, 2257–2263 (2016).

37. Ding, T. & Lew, M. D. Single-Molecule Localization Microscopy of 3D Orientation and Anisotropic Wobble Using a Polarized Vortex Point Spread Function. J. Phys. Chem. B 125, 12718– 12729 (2021).

38. Novotny, L. & Hecht, B. Principles of Nano-Optics. (Cambridge University Press, 2012).

39. Okuta, R., Unno, Y., Nishino, D., Hido, S. & Loomis, C. CuPy: A NumPy-Compatible Library for NVIDIA GPU Calculations. in Proceedings of Workshop on Machine Learning Systems (LearningSys) in The Thirty-first Annual Conference on Neural Information Processing Systems (NIPS) (2017).

40. Chiu, C.-L., Clack, N., & the napari community. napari: a Python Multi-Dimensional Image Viewer Platform for the Research Community. Microsc. Microanal. 28, 1576–1577 (2022).

41. Walt, S. van der et al. scikit-image: image processing in Python. PeerJ 2, e453 (2014).

42. Harris, C. R. et al. Array programming with NumPy. Nature 585, 357–362 (2020).

43. Tabaei, S. R. et al. Biomembrane Fabrication by the Solvent-assisted Lipid Bilayer (SALB) Method. J. Vis. Exp. JoVE 53073 (2015) doi:10.3791/53073.

44. Ferhan, A. R. et al. Solvent-assisted preparation of supported lipid bilayers. Nat. Protoc. 14, 2091–2118 (2019).

45. Ray, S., Mukherjee, K. & Bhattacharyya, S. N. Protocol to study internalization and localization dynamics of exogenously added proteins in detergent-permeabilized human cells. STAR Protoc. 6, 103992 (2025).

46. Boothe, T. et al. A tunable refractive index matching medium for live imaging cells, tissues and model organisms. eLife 6, e27240 (2017).

47. Mastronarde, D. N. Automated electron microscope tomography using robust prediction of specimen movements. J. Struct. Biol. 152, 36–51 (2005).

48. Gaifas, L., Kirchner, M. A., Timmins, J. & Gutsche, I. Blik is an extensible 3D visualisation tool for the annotation and analysis of cryo-electron tomography data. PLOS Biol. 22, e3002447 (2024).

49. Kremer, J. R., Mastronarde, D. N. & McIntosh, J. R. Computer visualization of three-dimensional image data using IMOD. J. Struct. Biol. 116, 71–76 (1996).

50. Liu, Y.-T. et al. IsoNet2 determines cellular structures at submolecular resolution without averaging. Preprint at 10.64898/2025.12.09.693325 (2025).

51. Lamm, L. et al. MemBrain: A deep learning-aided pipeline for detection of membrane proteins in Cryo-electron tomograms. Comput. Methods Programs Biomed. 224, 106990 (2022).

52. So-Last, M. G. F., Burt, A., Hale, T. & Allegretti, M. Easymode: general pretrained networks for cellular cryo-ET enable flexible approaches to subtomogram averaging. 2026.05.19.726344 Preprint at 10.64898/2026.05.19.726344 (2026).

