## Extended_Data for "Probing intracellular physical environments by rotational and translational single-particle tracking"

[1]Max Planck Insitute of Biophysics,
Group of Structure and Dynamics of Chromosomes,
60438 Frankfurt am Main, Germany

[2]Max Planck Insitute of Biophysics,
Group of Mechanisms of Cellular Quality Control,
60438 Frankfurt am Main, Germany

[3]University of Würzburg,
Nano-Optics and Bio-Photonics,
97074 Würzburg, Germany

### 1 Extended Data: Video Captions

1. Supplementary Video 1. AuNR tethered to a DOPC-bilayer at 60% glycerol concentration. The AuNR can be seen to diffuse and rotate freely on the DOPC-bilayer.

2. Supplementary Video 2. Comparison of intracellular AuNRs after delivery: cytoplasmic (left), endosomal fed (center) and endosomal starving (right).

### 2 Extended Data: Figures

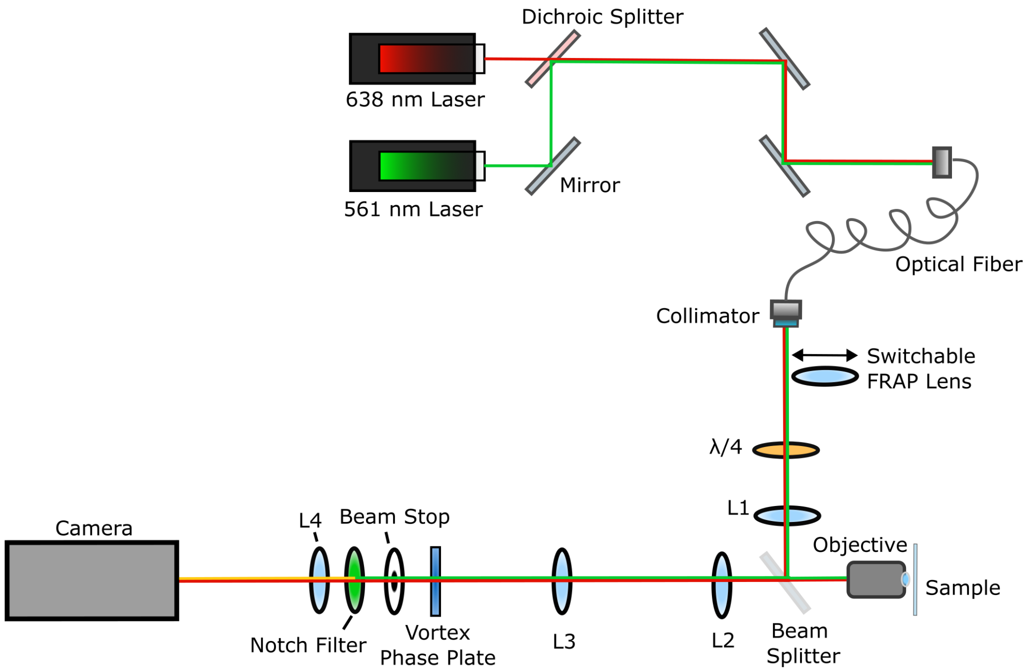


Supplementary Figure 1. Schematic of the optical setup for simultaneous rotational and translational single-particle tracking. Two excitation lasers (Cobolt 06-DPL 561 nm and Cobolt 06-MLD 638 nm) are coupled into a single optical fiber, with the output collimated at the fiber exit. A $\lambda/4$ wave plate generates circularly polarized illumination. A Köhler lens (L1; f = 150 mm) is positioned such that its focal plane coincides with the back focal plane of the objective (Zeiss Alpha Plan-Apochromat 100×/1.46 Oil), providing wide-field illumination. The excitation beam is reflected by a 50:50 beamsplitter into the objective. Scattered or fluorescent light collected from the sample is transmitted through the same beamsplitter and relayed by a 4f lens system (L2 and L3; f = 165 mm), which creates a conjugate plane of the objective back focal plane for placement of the vortex phase plate. A 5 mm beam stop is positioned immediately behind the vortex phase plate with a slight offset from the focal point of the reflected laser beam to generate a near-darkfield configuration that suppresses background signals. For fluorescence imaging, a 561 nm notch filter can be inserted before the final imaging lens (L4; f = 300 mm). A removable FRAP lens (f = -50 mm), placed before L1, enables localized photobleaching experiments. Images were acquired using a PCO Edge 4.2 camera.

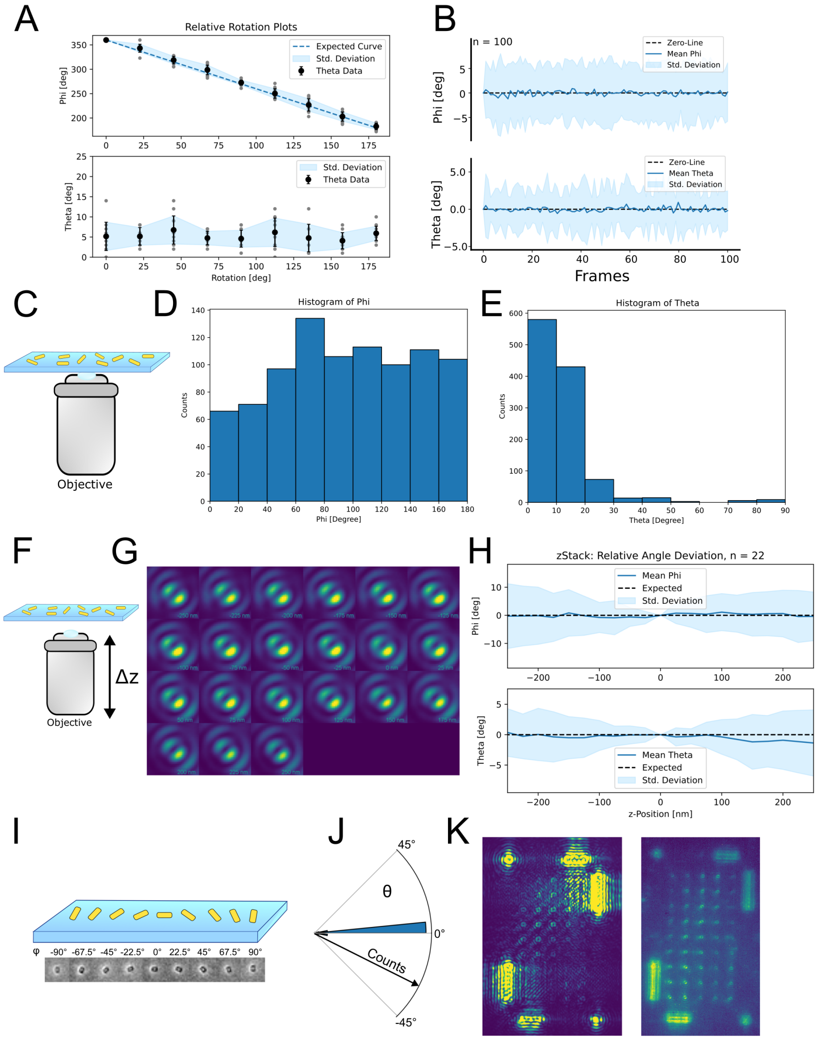


Supplementary Figure 2. Validation of rotational tracking accuracy and orientation measurements. a, Manual rotation of an AuNR in the azimuthal angle ($\phi$) and the corresponding detected polar angle ($\theta$) orientations (n = 18). b, Orientation stability of 100 surface-immobilized AuNRs measured over 100 consecutive frames, showing the standard deviation of the detected $\phi$ and $\theta$ angles relative to the reference orientation. c, Schematic of AuNRs randomly immobilized on a glass coverslip. d, The corresponding histogram of detected azimuthal angles ($\phi$) and e. the polar angles ($\theta$), n=1,000. f, Schematic of z-stack acquisition. g, Representative montage of an immobilized AuNR imaged over a z-stack spanning ±250 nm. h, Propagation of orientation errors in $\phi$ and $\theta$ as a function of defocus from the focal plane. i, Schematic and electron microscopy image of the nanofabricated AuNR grid. j, Polar histogram of detected $\theta$ orientations for the nanofabricated AuNR grid (n = 12). k, Representative images of the nanofabricated AuNR grid acquired in scattering and luminescence imaging modes.

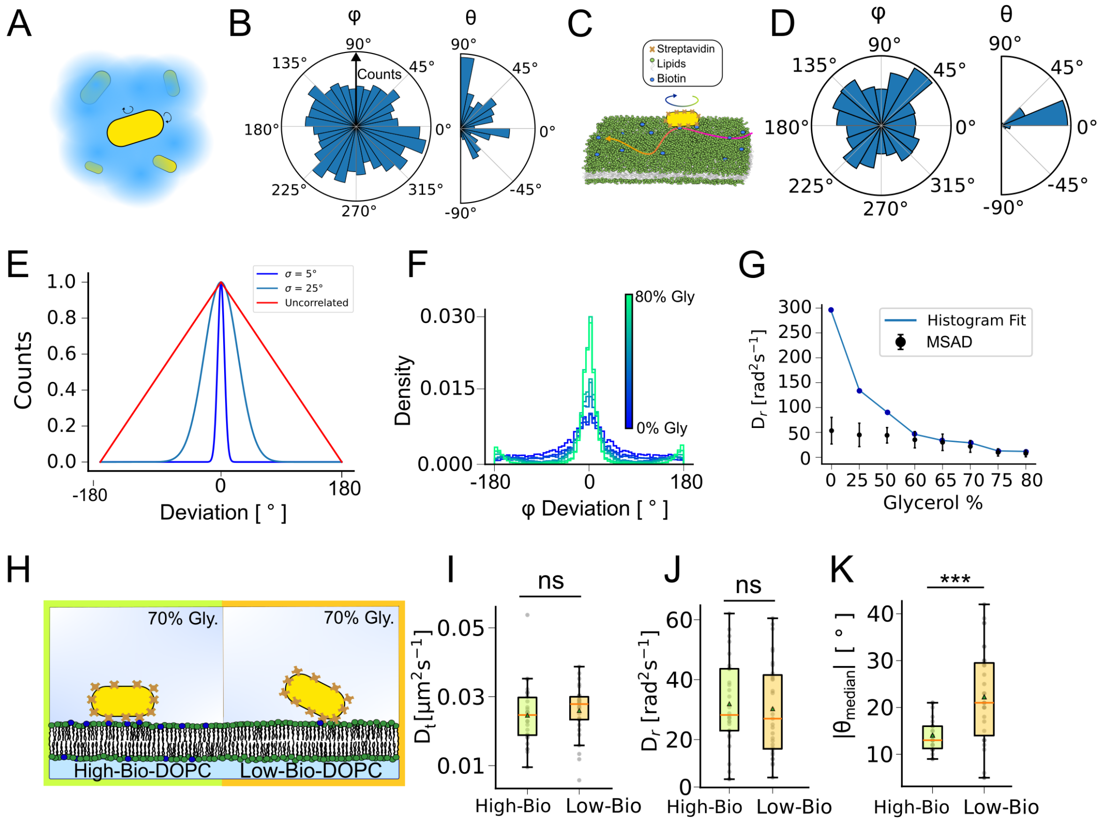


Supplementary Figure 3. Validation and characterization of rotational diffusion measurements. a, Schematic and b, Polar histograms of the azimuthal ($\phi$) and polar ($\theta$) orientations detected for freely diffusing AuNRs in 80% glycerol (n = 30). c, Schematic and d, Polar histograms of the detected $\phi$ and $\theta$ orientations for AuNRs tethered to a DOPC bilayer in an 80% glycerol–water mixture (n = 30). e, Simulated angular deviation distributions at different sampling frequencies for rotational diffusion with standard deviations ($\sigma$) of 5° and 25°, together with the triangular distribution expected for uncorrelated orientations. f, Angular deviation distributions of $\phi$ measured for AuNRs tethered to a DOPC bilayer over a glycerol titration (0%, 25%, 50%, 60%, 65%, 70%, 75%, and 80% glycerol). g, Comparison of rotational diffusion coefficients (Dr) obtained by mean squared angular displacement (MSAD) analysis and angular histogram fitting across the glycerol titration, showing convergence of the two methods at glycerol concentrations of $\geq$60%. h, Schematic of AuNR tethering to DOPC bilayers containing high (1:$10^3$) or low (1:$10^6$) biotin densities. i, Boxplots of translational diffusion and j. rotational diffusion coefficients for AuNRs tethered to high- and low-biotin DOPC bilayers, showing no significant differences (ns). k, Standard deviation of the unwrapped $\theta$ angle for AuNRs tethered to high- and low-biotin DOPC bilayers. Statistical significance was assessed using a two-tailed t-test (***, P $<$ 0.001).

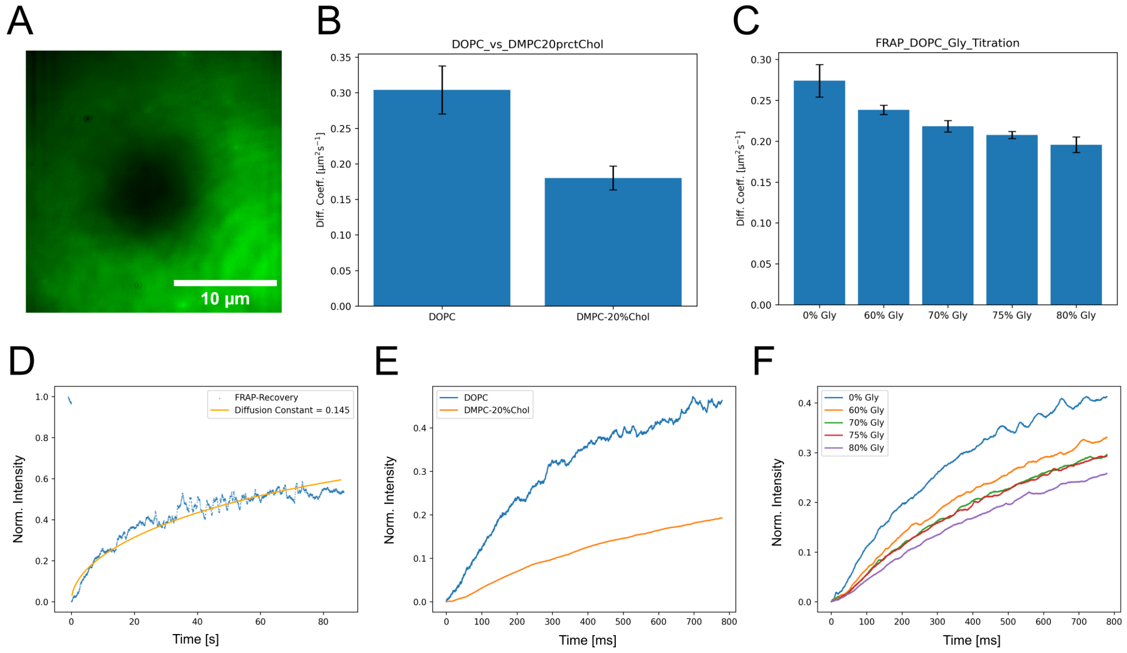


Supplementary Figure 4. Validation of membrane fluidity by FRAP. a, Representative image of the photobleached region on a DOPC bilayer during fluorescence recovery. b, Lateral diffusion coefficients of DOPC and DMPC + 20% cholesterol bilayers determined by FRAP. c, Lateral diffusion coefficients of DOPC bilayers measured over a glycerol titration by FRAP. d, Representative FRAP recovery curve (blue) and corresponding fit using the Soumpasis model (orange). e, Average FRAP recovery curves for DOPC (blue) and DMPC + 20% cholesterol (orange) bilayers. f, Average FRAP recovery curves for the DOPC glycerol titration (0%, 60%, 70%, 75%, and 80% glycerol).

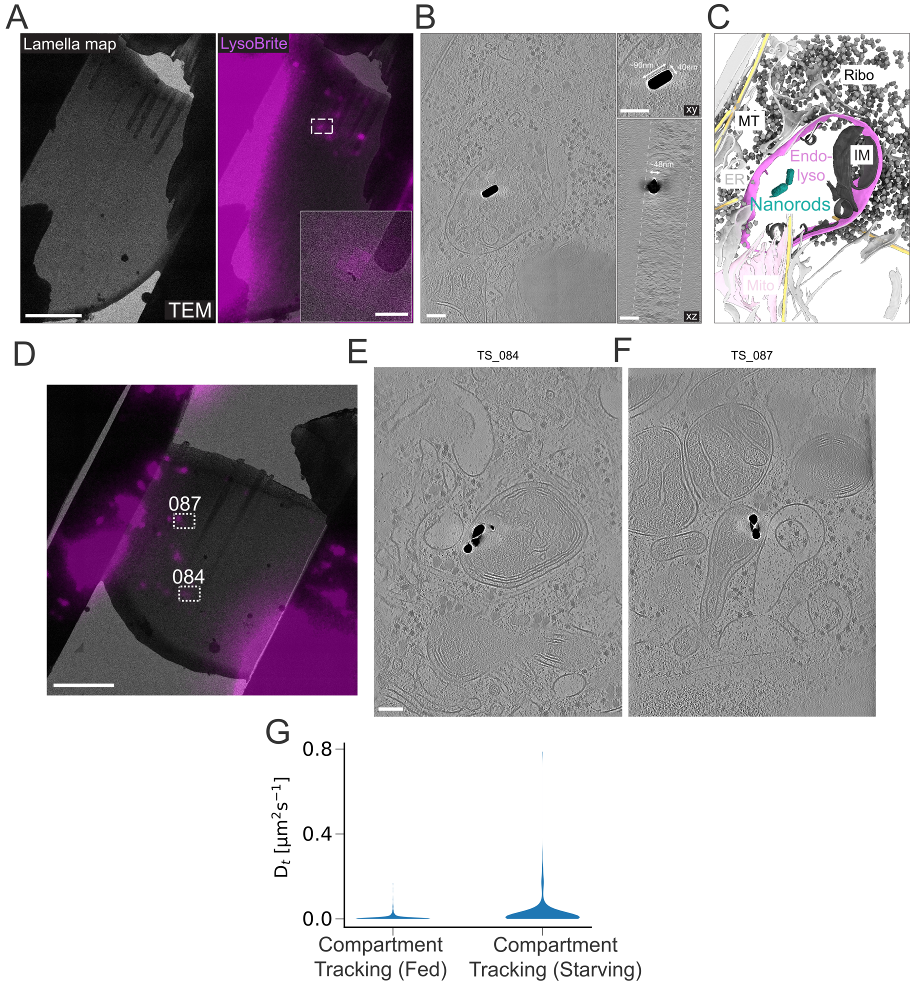


Supplementary Figure 5. Correlative cryo-fluorescence and cryo-electron tomography of intracellular AuNRs. a, Cryo-electron tomogram image of a FIB-milled lamella with post-milling LysoBrite fluorescence overlaid. The dashed rectangle indicates the tilt-series acquisition area, shown enlarged at right. Scale bars, 5 µm (lamella map) and 0.5 µm (zoom-in). b, Denoised cryo-ET tomographic slice showing an AuNR (dark) enclosed within a lysosomal compartment. Insets show enlarged views and AuNR dimensions measured in the xy and xz projections. Lamella boundaries are indicated by dashed lines in the xz view. Scale bar, 100 nm. c, Segmentation of the tomogram shown in b. Identified structures include the endo-lysosomal compartment (Endolyso), internalized membranes (IM), mitochondria (Mito), endoplasmic reticulum (ER), microtubules (MT), and ribosomes (Ribo). d, Second example of a FIB-milled lamella with post-milling LysoBrite fluorescence overlaid. The dashed rectangle indicates the tilt-series acquisition area. e,f, Denoised cryo-ET tomographic slices showing an AuNR enclosed within a lysosomal compartment. Scale bars, 5 µm (lamella map) and 100 nm (tomograms). g, Violin plots of translational diffusion coefficients ($D_t$) for LysoBrite-positive compartments in fed and starved cells.
