## Supplementary_Info for "Probing intracellular physical environments by rotational and translational single-particle tracking"

[1]Max Planck Insitute of Biophysics,
Group of Structure and Dynamics of Chromosomes,
60438 Frankfurt am Main, Germany

[2]Max Planck Insitute of Biophysics,
Group of Mechanisms of Cellular Quality Control,
60438 Frankfurt am Main, Germany

[3]University of Würzburg,
Nano-Optics and Bio-Photonics,
97074 Würzburg, Germany

**5 Theoretical prediction of free diffusion in glycerol** **12**

### 1 PSF Simulation

This section showcases the equations used in the Python-script to simulate PSFs from a scattering gold nanorod (AuNR) using full vectorial ray-tracing [1, 2].

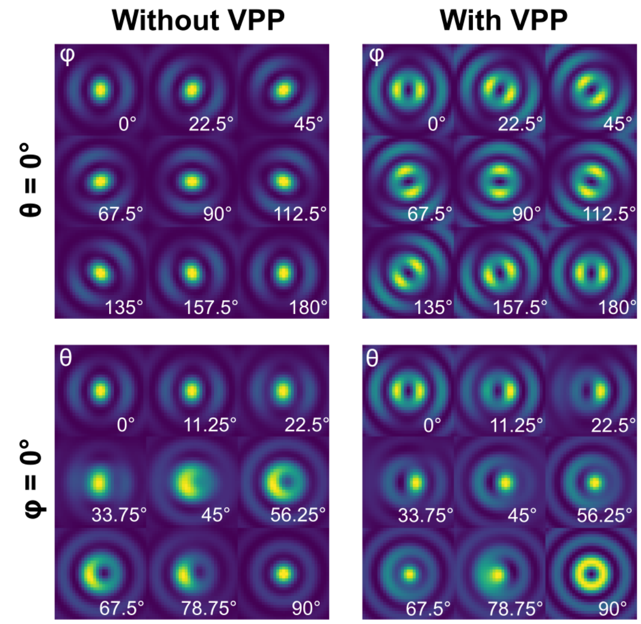

Figure 1: Exemplary intensity normalized simulated PSF rotations without and with VPP along $\phi$ and $\theta$, respectively.

#### 1.1 Equations

We define rotation matrices by an angle $\beta$ around each of the three axes,

$$M_{rotx}(\beta) = \begin{pmatrix} 1 & 0 & 0 \\ 0 & cos(\beta) & sin(\beta) \\ 0 & -sin(\beta) & cos(\beta) \end{pmatrix} \tag{1}$$

$$M_{roty}(\beta) = \begin{pmatrix} cos(\beta) & 0 & sin(\beta) \\ 0 & 1 & 0 \\ -sin(\beta) & 0 & cos(\beta) \end{pmatrix} \tag{2}$$

$$M_{rotz}(\beta) = \begin{pmatrix} cos(\beta) & sin(\beta) & 0 \\ -sin(\beta) & cos(\beta) & 0 \\ 0 & 0 & 1 \end{pmatrix} \tag{3}$$

where clockwise rotation around $\beta$ can be achieved either by a negative $\beta$ or by transposing the matrix.

From the refractive index $n$ and absorption index $\kappa$ retrieved from Johnson & Christy [3], we use following relationships to obtain $\epsilon_1$ and $\epsilon_2$,

$$\epsilon_1 = n^2 - \kappa^2, \epsilon_2 = 2n\kappa \tag{4}$$

and hence the complex relative permittivity value

$$\epsilon = \epsilon_1 + \epsilon_2 \cdot i. \tag{5}$$

We define the polarizability tensor $\alpha$ for a prolate ellipsoid according to Bohren & Hoffman [4] with the diameter $d$, the aspect ratio $A$, the permittivity of gold $\epsilon_g$ and the permittivity of the medium $\epsilon_m$.

With the eccentricity $e_c$ defined as:

$$e_c = \sqrt{1 - (\frac{1}{A})^2}. \tag{6}$$

We then define $l_1$ as

$$l_1 = (\frac{1 - e_c^2}{e_c^2}) \cdot (\frac{1}{2} ecc \log(\frac{1 + e_c}{1 - e_c}) - 1) \tag{7}$$

and $l_2$ as

$$l_2 = \frac{1 - l_1}{2} \tag{8}$$

allowing us to obtain the polarizability along the AuNR's long axis

$$\alpha_1 = 4\pi A(\frac{d}{2})^3 \cdot (\frac{\epsilon_g - \epsilon_m}{3\epsilon_m + 3l_1(\epsilon_g - \epsilon_m)}) \tag{9}$$

and the polarizability along the AuNR's short axis

$$\alpha_2 = 4\pi A(\frac{d}{2})^3 \cdot (\frac{\epsilon_g - \epsilon_m}{3\epsilon_m + 3l_2(\epsilon_g - \epsilon_m)}) \tag{10}$$

resulting in the polarizability-tensor:

$$\alpha = \begin{pmatrix} \alpha_1 & 0 & 0 \\ 0 & \alpha_2 & 0 \\ 0 & 0 & \alpha_2 \end{pmatrix} \tag{11}$$

From this we can obtain the electric Dipole-moment $\mathbf{p}$ of a AuNR driven by an incident electric field $\mathbf{E_i n}$ (we use left-handed circularly polarized light) as

$$\mathbf{p} = \mathbf{E_{in}} \cdot \alpha. \tag{12}$$

We then provide the ray-tracing starting vectors $\mathbf{k_n}$ for a scatterer as functions of direction defined by the angles $\theta$ and $\phi$:

$$\mathbf{k_n}(\theta,\phi) = M_{rotz}(-\phi)\cdot\begin{pmatrix} sin(\theta)\\ 0\\ cos(\theta)\end{pmatrix}, \tag{13}$$

whereas the corresponding field emitted by the dipole in that direction:

$$\mathbf{E}_{dipole}(\theta,\phi) = (\mathbf{k_n}(\theta,\phi)\otimes\mathbf{p})\otimes\mathbf{k_n}(\theta,\phi) \tag{14}$$

Next, we define a ray-tracing lens-matrix $L(x)$ for an aplanatic lens with the apodization factor $x$. It fulfills the condition $x = 1$ for focusing and $x = 2$ for collimation.

$$L(x,\beta) = \begin{cases} M_{roty}(\beta) & \text{, if } x = 0\\ \sqrt{cos(\beta)}\cdot M_{roty}(\beta) & \text{, if } x = 1\\ \frac{1}{\sqrt{cos(\beta)}}\cdot M_{roty}(\beta) & \text{, if } x = 2\end{cases} \tag{15}$$

We further define functions to describe a vortex phase-plate (VPP) of order m (we use $m = 1$ )

$$\sigma_{VPP}(\phi) = e^{i\cdot m\cdot\phi} \tag{16}$$

and the beamstop (BS), as a function of the angle $\theta$ and the $\theta$-threshold $T$ (in our case $T = 53°$):

$$\sigma_{BS}(\theta) = \begin{cases} 10^{-3} & \text{, if } \theta < T\\ 1 & , \; else\end{cases} \tag{17}$$

Moreover we obtain the angle at the detector plane $\theta_2 = \arcsin(\frac{\theta}{M})$,where $M$ is the Magnification of our optical system (in our case $M = 180$) .

With these components, we can obtain field-components $\mathbf{E}_{det}(\theta,\phi)$ for the emission directions *theta* and *phi* at the detector plane

$$\mathbf{E}_{det}(\theta,\phi) = [M_{rotz}(-\phi)\cdot L(1,-\theta_2)\cdot L(2,-\theta)\cdot M_{rotz}(\phi)\cdot\mathbf{E}_{dipole}(\theta,\phi)]\cdot\sigma_{VPP}(\phi)\cdot\sigma_{BS}(\theta), \tag{18}$$

as well as the corresponding direction vectors:

$$\mathbf{k}_{det}(\theta,\phi) = M_{rotz}(-\phi)\cdot L(0,-\theta_2)\cdot L(0,-\theta)\cdot M_{rotz}(\phi)\cdot\mathbf{k}_n(\phi,\theta) \tag{19}$$

Finally, the total electric field at the detector locations r = $(x, y, 0)$ can be obtained as

$$\mathbf{E}(\mathbf{r}) = \sum_{i,j}\mathbf{E}_{det}(\theta_i,\phi_j)\cdot\exp\left(\frac{2\pi}{\lambda}i\cdot(\mathbf{r}\cdot\mathbf{k}_{det}(\theta_i,\phi_j)\right), \tag{20}$$

where $\lambda$ is the wavelength and $\theta_i$ and $\phi_j$ are a defined set of equally spaced angles ranging from 0 to arcsin(0.95) (max collection angle) and 0 to $2\pi$ respectively (we used 32 angles for each range).

The corresponding intensities are then obtained using $I \propto |\mathbf{E}|^2$.

#### 2 Brute-Force-Fitting Routine

To fit the raw data to corresponding orientations, we devised a fast, GPU-aided Brute-Force-Fitting (BFF) routine in Python using CuPy [5] and Napari [6]. Because computational resources and time are essential for quick experimental feedback, our custom Python software allows the adjustment of parameters to achieve a balance between computational expense and goodness of fit, especially when dealing with very large amounts of data. The adjustment of parameters for the routine also allowed us to adapt the analysis to the versatile raw data that was recorded.

Beforehand, we simulate a full rotation covering all angle combinations in 1° steps. This simulation is then subsequently used for the BFF-subtraction, which avoids the need to compute orientations for every iteration. To be able to fit the raw data successfully, the measured PSFs have to be extracted and properly centered into an array of the same size as the simulation. For more information, please read section 4.1 about PSF extraction.

Generally, the BFF method relies on subtracting the raw frame from the simulated stack, resulting in residuals for each subtracted position in the stack. Then, the least squares method is used to evaluate which position in the stack is a match for the raw detected PSF. To evaluate the goodness of the fit, an R-squared value is calculated for each subtracted position in the simulation. A summary sketch of this basic pipeline can be seen on Fig. 2.

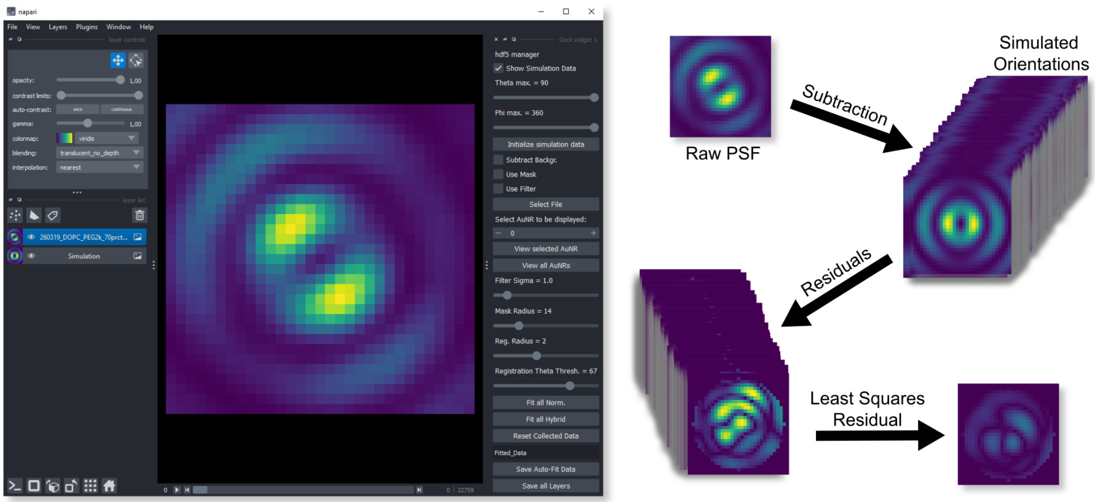

Figure 2: A snapshot of the BFF-Program GUI with a sketch demonstrating the basic pipeline.

Furthermore, the program allows the user to do basic pre-processing of the raw data: a gaussian filter with adjustable kernel and simple background subtraction which subtracts the minimum value of the background to the whole image. This will eliminate spurious noise and level the raw data's background closer to zero, which avoids excessive residuals. Adding a circular mask to the subtraction and adjusting its radius is optional and can optimize fitting conditions and results.

The pre-centering of AuNRs for fitting exhibits an intrinsic error margin, or wobble, that will vary throughout the measured stack around a few pixels. To counteract the effect of poorly centered

PSFs, we incorporate an image registration routine for the subtraction, along a square around the center position. The position of the square is determined by the registration radius in pixel, and for each position in the registration, a new subtraction is performed. This allows us to compare residuals along the image registration and choose the residual with global least squares. With increasing radius, the amount of subtractions will increase exponentially because the registration is performed along a square of pixels with the length $2r + 1$, with r being the radius and the center pixel being added for symmetricity.

Wobbling of the pre-centered PSFs will be reduced at high signal orientations, therefore, at low or flat $\theta$ angles it will increase. To reduce computational resources for well-centered orientations, a threshold can optionally be set for theta to start a registration if the initial fit exceeds that value in $\theta$. This will also help minimize false high angle detections that might come from misaligned PSFs.

The program offers two fit-modes, hybrid-fitting and normalized-fitting. Hybrid fitting is recommended for most use-cases and normalized fitting is to be used with caution, due to its propensity for higher frequency of false orientation detection under certain conditions. The following two sub-chapters explain these approaches in-depth.

#### 2.1 Hybrid-BFF

AuNRs will exhibit progressively lower signal when their orientation in $\theta$ increases (Fig. 3). This intrinsic information encoded in the detected intensity can be used to aid the BFF-results for higher $\theta$ angles, as they can be particularly difficult to fit correctly.

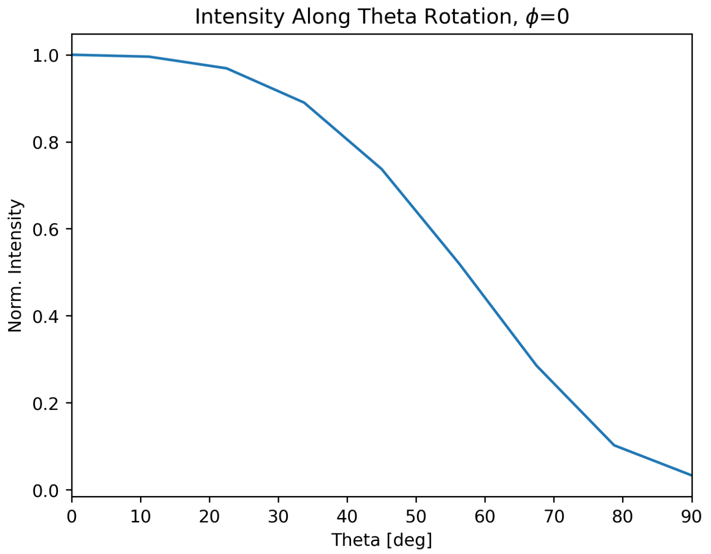

Figure 3: Decreasing average normalized intensity for a simulated rotation along $\theta$ for $\phi = 0$.

The hybrid fit uses two sets each of differently normalized simulated data and acquired PSFs. The first normalization happens frame-to-frame, where each frame is normalized to its maximum value. This eliminates the intensity fluctuations along the datasets. Secondly, the data is normalized to the maximum intensity slice in the stack. This will normalize the stack to the highest intensity slice, both for the simulated and acquired data. This allows to incorporate intensity information into the BFF-subtraction to determine $\theta$ orientations more robustly. However, this will only work, if it is assumed that the experimentally acquired PSFs also exhibit flat orientations or maximum intensity slices in their stacks.

With these sets of differently normalized data, the routine will first subtract the frame-to-frame normalizations. If a $\theta$ orientation is detected above a set threshold, the fit will re-occur with the datasets normalized to the maximum slice, containing the intensity information. Additionally, an image registration routine will be performed, as discussed above.

#### 2.2 Normalized-BFF

The normalized BFF happens on a frame-top-frame normalization basis for both the simulation and the acquired data. Because it does not take into account intensity information or intensity fluctuations along the data, it is more prone to errors for high $\theta$ orientations and therefore only recommended for very narrow applications, like completely flat AuNRs.

### 3 Bilayer FRAP

Fluorescence Recovery after Photobleaching (FRAP) experiments were performed on lipid bilayers to confirm our observations from translational diffusion of AuNRs. To allow localized and focused FRAP experiments with a diameter of approximately 10 µm, an additional -50 mm focal length lens was placed before the Koehler lens of the setup. Before photobleaching, a baseline was recorded at low laser power (10 mW), followed by a bleaching procedure at the maximum laser power of 120 mW. The recovery was then continuously monitored at 10 mW laser power for the rest of the acquisition. For the experiments, a 561 nm laser was used. The bilayer was labeled with Liss-Rhodamine-PE at a concentration of 0.25 mol% (Avanti Research).

The FRAP curve was double normalized to its minimum and maximum value, where the maximum value is taken from the reference baseline before photobleaching. Subsequently, the curve was fit with the Soumpasis model as follows [7]:

$$I(t) = e^{\frac{-2\tau}{t}}(J_0(\frac{2\tau}{t}) + J_1(\frac{2\tau}{t})) \tag{21}$$

With $J_i$ being the modified Bessel function of order i and $\tau$ being defined as $\tau = \frac{w^2}{4D}$ with width $w$ and diffusion coefficient $D$.

### 4 Analysis Pipeline

#### 4.1 PSF- and centroid-extraction

To allow proper fitting of the orientations exhibited by individual AuNRs throughout the acquired stacks, PSFs have to be extracted from the raw data and centered in a 31x31 pixel array to match the simulations. Additionally, throughout the PSF extraction procedure, the global centroids are extracted as well, to reconstruct translational trajectories. This sub-section elucidates these procedures.

We developed a custom Napari-based [6] Python software and pipeline for automatic PSF and centroid extraction. The pipeline starts with detection of the AuNR in the first slice. Around this first localization, a region-of-interest (ROI) is cropped for further processing. The extraction pipeline continues with a TopHat filter, followed by a gaussian filter and an inverted Sato filter from Scikit-Image [8], which will take advantage of the minima of the diffraction rings around the PSF to outline its position, and most importantly, the dark spot inside the PSF will become foreground.

Then a simple threshold $> M + x \cdot \sigma$ is applied to the Sato-filtered ROI, where $M$ is the median, $x$ is a factor determined by the user and $\sigma$ is the standard deviation of the ROI. The resulting threshold binary image is used as a mask on the Sato-filtered ROI. Finally, the resulting regions in the masked image are analysed separately for their intensity content, and the maximum intensity region's centroid is used to crop a 31x31 pixel area, which centers the PSF. The global centroid is saved as the coordinate of the AuNR. An example of the pipeline can be seen on Fig. 4.

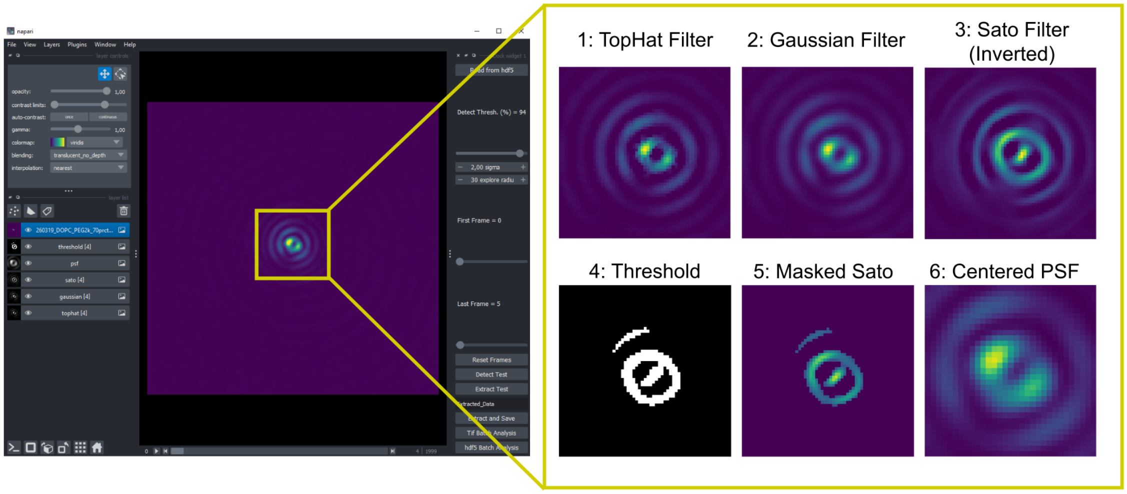

Figure 4: Snapshot of the custom Napari-based PSF-extraction software with an example extraction routine outlined for a single frame.

After extracting and centering the PSF the program continues to the next frame, where a search radius given by the user will determine how far away from the previous localization the software will look for the next localization. If a new localization is found inside this search radius, the routine is repeated for the new localization in the current frame. Following this search procedure for each frame, coupled with the outlined processing pipeline, the software will automatically extract the PSF at all given positions during the trajectory and reconstruct its lateral coordinates as well.

#### 4.2 Angle unwrapping and rotational trajectories

Angles were unwrapped using Python's Numpy function numpy.unwrap [9]. The period limit for $\phi$ was set to be 360° with a discont of 180°, as the maximum resolution in $\phi$ is 180° because of angle degeneracy above this threshold. The period selected for theta was 180° with a discont of 90°.

Rotational trajectories were reconstructed from the fit orientations by defining one of the AuNR termini along the long-axis as the reference point. The trajectory is always reconstructed following the condition of the shortest path traversed by the reference terminus. This allows the trajectory to also explore negative angles in $\theta$, even though the BFF fitting routine only outputs $\theta$ values from 0° to 90°. To incorporate negative $\theta$ orientations, the following condition was used, where $T_\phi$ is the period threshold for unwrapping in $\phi$:

$$\theta_{new} = \theta \cdot n \begin{cases} n = 1 & \text{, if } \Delta\phi < T_\phi \\ n = -1 & \text{, if } \Delta\phi > T_\phi \end{cases} \tag{22}$$

In other words, every time a step larger than $T_\phi$ is unwrapped, $\theta$ turns negative. This fulfills the above stated condition where we trace the shortest path traversed by the reference AuNR terminus.

#### 4.3 MSD/MSAD Analysis

Mean squared displacement (MSD) and Mean squared angular displacement (MSAD) analysis was performed following the general derivation for MSD for n dimensions, as follows:

$$MSD = \langle (x_1(t) - x_1(0))^2 \rangle + \langle (x_2(t) - x_2(0))^2 \rangle + ... + \langle (x_n(t) - x_n(0))^2 \rangle \tag{23}$$

with $MSD = 2nDt$, where $t$ is time and $D$ represents the diffusion coefficient. The special cases for the translational MSD and the rotational MSAD results in following two equations:

$$MSD = \langle (x(t) - x(0))^2 \rangle + \langle (y(t) - y(0))^2 \rangle + \langle (z(t) - z(0))^2 \rangle \tag{24}$$

for the translational MSD, and

$$MSAD = \langle (\phi(t) - \phi(0))^2 \rangle + \langle (\theta(t) - \theta(0))^2 \rangle \tag{25}$$

for rotational MSAD.

The MSD/MSAD was then fit with the following power-law model:

$$Fit = K \cdot x^\alpha \tag{26}$$

with $K$ being the slope of the power-law and $\alpha$ the exponent. The exponent gives insight into the diffusion behavior as follows:

$$\alpha \begin{cases} < 1\text{, subdiffusion} \\ = 1\text{, brownian motion} \\ > 1\text{, active diffusion} \end{cases} \tag{27}$$

and the slope relates back to the diffusion coefficient with the relationship $K = 2nD$, where $n$ is the number of independent dimensions. An example MSAD fit can be seen on Fig. 5.

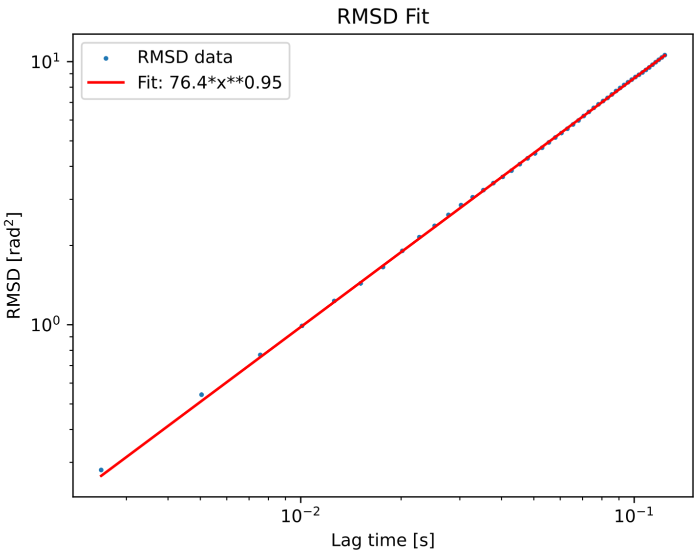

Figure 5: Example MSAD fit using a power-law model.

### 5 Theoretical prediction of free diffusion in glycerol

Following Tirado et al. [10], the following equation was used to determine the theoretical rotational diffusion coefficients $D_r$ for a freely diffusing, flat-ended cylinder in a given viscosity:

$$D_r = \frac{3k_B T}{\pi \eta L^3}(ln(a) + \delta_\perp) \tag{28}$$

where $T$ is the temperature, $k_B$ is the Boltzmann constant, $\eta$ is the viscosity, $L$ is the length of the AuNR and $\delta_\perp$ is the correction $\delta_\perp = -0.662 + \frac{0.917}{a} - \frac{0.050}{a^2}$ with $a = \frac{L}{d}$, with d being the AuNR's diameter.

Because the population of AuNRs is never identical in a colloidal solution, we consulted the diameter deviation specified by the supplier of our AuNRs (Nanopartz), which corresponded to ± 5 nm for this specific batch. According to this deviation, we calculated the theoretical lower and upper range of $D_r$, as well as the theoretical $D_r$ for the correctly sized AuNRs. The diameter of the AuNRs used in the experiments was 40 nm with an aspect ratio of 2.

| Glycerol % | 80% | 85% | 90% | 95% | 99% |
|---|---|---|---|---|---|
| Theoretical upper $D_r$ [$rad^2 s^{-1}$] | 82.5 | 48.3 | 26.4 | 13.3 | 8.4 |
| Theoretical $D_r$ [$rad^2 s^{-1}$] | 55.3 | 32.3 | 17.7 | 8.9 | 5.6 |
| Theoretical lower $D_r$ [$rad^2 s^{-1}$] | 38.8 | 22.7 | 12.4 | 6.2 | 3.9 |
| Experimental mean $D_r$ [$rad^2 s^{-1}$] | 50.0 | 37.4 | 23.1 | 19.5 | 11.1 |

Table 1: Theoretical and experimental rotational diffusion coefficients.

#### References

[1] Lukas Novotny and Bert Hecht. *Principles of nano-optics.* Cambridge university press, 2012.

[2] Jeongmin Kim, Yuan Wang, and Xiang Zhang. Calculation of vectorial diffraction in optical systems. *Journal of the Optical Society of America A*, 35(4):526, 2018.

[3] Peter B Johnson and R-WJPrB Christy. Optical constants of the noble metals. *Physical review B*, 6(12):4370, 1972.

[4] Craig F Bohren and Donald R Huffman. *Absorption and scattering of light by small particles.* John Wiley & Sons, 2008.

[5] Ryosuke Okuta, Yuya Unno, Daisuke Nishino, Shohei Hido, and Crissman Loomis. CuPy: A NumPy-Compatible Library for NVIDIA GPU Calculations. In *Proceedings of Workshop on Machine Learning Systems (LearningSys) in The Thirty-first Annual Conference on Neural Information Processing Systems (NIPS)*, 2017.

[6] Chi-Li Chiu, Nathan Clack, and the napari community. napari: a Python Multi-Dimensional Image Viewer Platform for the Research Community. *Microscopy and Microanalysis*, 28(S1):1576–1577, August 2022.

[7] DM1329018 Soumpasis. Theoretical analysis of fluorescence photobleaching recovery experiments. *Biophysical journal*, 41(1):95–97, 1983.

[8] Stéfan van der Walt, Johannes L. Schönberger, Juan Nunez-Iglesias, François Boulogne, Joshua D. Warner, Neil Yager, Emmanuelle Gouillart, and Tony Yu. scikit-image: image processing in Python. *PeerJ*, 2:e453, June 2014.

[9] Charles R. Harris, K. Jarrod Millman, Stéfan J. van der Walt, Ralf Gommers, Pauli Virtanen, David Cournapeau, Eric Wieser, Julian Taylor, Sebastian Berg, Nathaniel J. Smith, Robert Kern, Matti Picus, Stephan Hoyer, Marten H. van Kerkwijk, Matthew Brett, Allan Haldane, Jaime Fernández del Río, Mark Wiebe, Pearu Peterson, Pierre Gérard-Marchant, Kevin Sheppard, Tyler Reddy, Warren Weckesser, Hameer Abbasi, Christoph Gohlke, and Travis E. Oliphant. Array programming with NumPy. *Nature*, 585(7825):357–362, September 2020.

[10] Maria M Tirado and José Garcia De La Torre. Rotational dynamics of rigid, symmetric top macromolecules. application to circular cylinders. *The Journal of Chemical Physics*, 73(4):1986–1993, 1980.
